# H2A.Z histone variants facilitate HDACi-dependent removal of H3.3K27M mutant protein in paediatric high-grade glioma cells

**DOI:** 10.1101/2023.05.15.540760

**Authors:** Katarzyna B. Leszczynska, Amanda Pereira de Freitas, Chinchu Jayaprakash, Monika Dzwigonska, Kamil Wojnicki, Bartlomiej Gielniewski, Paulina Szadkowska, Beata Kaza, Maciej K. Ciolkowski, Joanna Trubicka, Wieslawa Grajkowska, Bozena Kaminska, Jakub Mieczkowski

## Abstract

Diffuse intrinsic pontine gliomas (DIPG) are deadly paediatric brain tumours, non-resectable due to brainstem localisation and diffusive growth. Patients with DIPG have a dismal prognosis of 9-12 months of survival with no effective therapy. Over 80% of DIPGs harbour a mutation in histone 3 (H3.3 or H3.1) resulting in a lysine to methionine substitution (H3K27M). H3K27M causes global epigenetic alterations (a loss of H3K27 trimethylation and an increase in H3K27 acetylation) resulting in aberrant gene expression. To date, no therapeutic strategy exists to suppress the levels of oncogenic H3K27M.

We show that pan-HDAC inhibitors (HDACi) lead to the temporary but significant reduction in the H3.3K27M protein (up to 80%) in multiple glioma cell lines expressing the H3.3K27M histone variant, without changes in the *H3F3A* mRNA expression. The H3.3K27M occupancy at the chromatin is greatly reduced upon HDACi (SB939) treatment, as shown by ChIPseq analysis. H3.3K27M loss is most striking at SB939-upregulated genes suggesting the role in repression of these genes. In addition, genes previously reported as H3K27M-dependent become downregulated in response to SB939 treatment. We discover that the SB939-mediated loss of H3.3K27M is partially blocked by a lysosomal inhibitor, chloroquine. Moreover, the loss of H3.3K27M is facilitated by co-occurrence of H2A.Z, as evidenced by the knock-down of H2A.Z histone isoforms. ChIPseq analysis confirms the occupancy of H3.3K27M and H2A.Z at the same SB939-inducible genes.

Altogether, we provide new insight into disease-specific mechanism of HDAC inhibition and demonstrate pharmacological modulation of the oncogenic H3.3K27M protein levels. These findings open a new possibility to directly target the H3.3K27M oncohistone, which may be exploited in future therapies.

## INTRODUCTION

In eukaryotes, the chromatin structure is organised through nucleosomes composed of genomic DNA wrapped around cores histones ^1^. H2A, H2B, H3 and H4 are the four types of core histones, represented by ‘canonical’ replication-dependent histones, or replication-independent histone variants.^2^. The presence of histone variants in nucleosomes plays a particular role in regulating the chromatin structure and gene expression, and, as it is in case of H2A.Z and H3.3 histone variants, may have opposing effects ^3^. H2A.Z is concentrated at the promoters of inducible genes with low basal expression and is removed during transcription. ^4, 5^. Recent studies indicate that H2A.Z can even inhibit gene transcription ^6^. Conversely, H3.3 is regarded as a marker of actively transcribed genes ^7, 8^. The presence of particular histone variants proves critical in certain diseases, and particularly, in tumorigenesis.

Paediatric high-grade gliomas (pHGGs) of the midline brain structures such as pons, thalamus and spinal cord are classified as H3K27-altered diffuse midline gliomas (DMG), due to the global loss of lysine 27 trimethylation (H3K27me3) and a corresponding increase in H3K27 acetylation (H3K27ac) ^9–12^. In most cases, the H3K27 hypomethylation is caused by non-synonymous histone mutation resulting in a Lys27 (K27) to methionine (M) substitution mainly in H3.3, and to a lesser extent in H3.1 histones (H3K27M). In paediatric diffuse intrinsic pontine gliomas (DIPGs), the mutated H3K27M histone variant is expressed in more than 80% of cases ^9, 10^. H3K27M directly binds Enhancer of Zeste Homolog 2 (EZH2), the methyltransferase unit of the Polycomb Repressive Complex 2 (PRC2). This interaction inhibits the catalytic activity of PRC2, resulting in the loss of H3K27me3. PRC2 can also be inhibited by overexpression of EZH Inhibitory Protein (EZHIP), which occurs in K27-altered DMGs that do not express the H3K27M oncohistone ^9–11, 13–16^. Loss of H3K27me3 leads to altered activities of specific gene promoters and consequent transcriptional changes. The presence of the H3K27M drives expression of so called ‘H3K27M signature genes’, including PRC2 target genes, BMI1 and NOTCH pathway genes along with other genes involved in neurogenesis ^9, 17, 18^.

The H3K27M oncohistones coexist with other alterations, including the loss of p53 function, amplification of *PDGFA* or overexpression of *MYC/MYCN.* As shown with CRISPR and shRNA studies, removal of the mutated histone can reverse its oncogenic function, restore the distribution of H3K27me3 and H3K27ac modifications at the chromatin and normalise the expression of ‘H3K27M signatures gene’ ^19, 20^. However, currently there are no ways to eradicate this mutation from the tumours pharmacologically and numerous efforts had been made to target other epigenetic vulnerabilities of DMGs instead ^21^. Promising results in pre-clinical studies had been achieved with histone deacetylase inhibitors (HDACi), and in particular, with panobinostat, which induced glioma cell death, impaired the growth of tumour xenografts ^22^. Panobinostat also partially restored the levels of H3K27me3 in H3K27M-expressing DMG cells and reversed the expression of ‘H3K27M signatures’, although the mechanism behind this indirect action of the drug is unclear and had been speculated to rely on the poly-acetylation of the H3 N-terminal tail ^22–24^. In ensuing clinical trials, the use of panobinostat as a single agent did not improve the outcome of patients. Combinatorial treatments with other drugs, e.g. with proteasomal or transcriptional inhibitors showed promising effects encouraging clinical trials ^25, 26^. New therapeutics for children with DIPG are desperately needed as the current treatment relies only on radiotherapy, which in more than 90% of cases ends in disease recurrence and death within 9-12 months after diagnosis ^27^.

In this study, we show that pan-HDAC inhibitors (including pracinostat/SB939, panobinostat, vorinostat and entinostat) lead to a significant decrease in H3.3K27M protein levels, without affecting the *H3F3A* mRNA expression. The genome-wide analysis in SB939-treated cells confirmed the effects on H3K27M occupancy and dependent genes. We demonstrate that SB939-dependent loss of H3.3K27M is impaired by treatment with chloroquine, the inhibitor of autophagosome/lysosomal degradation and DNA intercalating agent. The loss of H3.3K27M protein upon SB939 treatment is facilitated by co-occurrence of non-canonical H2A.Z histone variants. Altogether, we show a new disease-specific mechanism of action of HDACi, which may pave ways to new therapeutic approach in these deadly paediatric tumours.

## RESULTS

We have performed a cell viability screen with selected drugs targeting epigenetic modifiers to search for compounds that could be efficient in targeting epigenetically-vulnerable pHGG cells expressing H3K27M. We initially used three cell lines originating from the paediatric brainstem HGG (SF8628, SF7761 and WG27) (**Figure 1A and Supplementary Table 1**). Panobinostat and GSK-J4 were included as a reference, as these previously showed effectiveness against pHGGs (**Figure 1A**) ^22, 28^. Several of the drug candidates led to a loss in cell viability at 10 µM concentrations (JIB-04, Methylstat, UNC0638, UNC0642, SB939/pracinostat) and were therefore further tested in a panel of cell lines and at a range of concentrations (**Figure 1B and Supplementary Figure 1A-E**). We found that all the shortlisted candidate drugs significantly decreased cell viability at sub-micromolar concentrations, but SB939, which is an HDACi, was less efficient against the WG27 cells, which expressed lower amounts of the mutated histone H3K27M when compared to other cell lines (**Figure 1A-B and Supplementary Figure 1F-G**). The effect of SB939 has not been previously characterised in H3K27M^+^ pHGGs; here we show that it induced apoptosis and decreased cell proliferation in these cells (**Supplementary Figure 1H-I**). We then further investigated whether the presence of H3.3K27M mutation confers increased sensitivity towards SB939. We examined cell viability in additional six cancer patient-derived pHGG cell cultures with different H3.3K27M status (**Figure 1C** and **Supplementary Table 1**). We observed that H3.3K27M^+^ cells were significantly more sensitive to SB939 treatment compared to cells with wild type H3.3 (approximately 50% of H3.3K27M^+^ cells survived 72 hour treatment with 1 µM SB939, while more than 50% of H3.3 wild type cells survived 72 hour treatment with 10 µM SB939, 2-way ANOVA *P=0.019,* **Figure 1C**). To further verify this observation, we used previously published isogenic pairs of cells, in which the H3.3K27M mutation was eliminated with CRISPR technology from the parental cell line (**Supplementary Table 1)** ^29, 30^. We observed a higher sensitivity to the drug at lower concentrations (two-way ANOVA *P=0.005*) when the H3.3K27M mutant was present only in one out of three tested pairs of cell lines (HSJ019) (**Figure 1D-E** and **Supplementary Figure 1J**). However, in all three tested pairs of cell lines (even in cells with deleted H3.3K27M by CRISPR), 50% cell killing occurred at concentrations below 1 µM SB939 (**Figure 1D-E and Supplementary Figure 1J)**. We also compared sensitivity of normal human astrocytes (NHA), which do not contain the H3 mutation, and these cells required higher SB939 concentrations (above 1 µM) to achieve 50% cell kill **(Supplementary Figure 1K)**. Together, these data suggested that pHGG cells that have the H3.3K27M mutation (or had it originally before gene deletion) are more sensitive to SB939 compared to H3 wild type pHGG cells. However, since the deletion of the mutation in the isogenic system did not conclusively decrease the sensitivity towards SB939 in all cell lines (**Figure 1D-F**), this suggests that other factors are also involved in the sensitivity to HDACi, which likely evolved during the growth of H3K27M^+^ pHGGs tumours.

**Figure 1.**
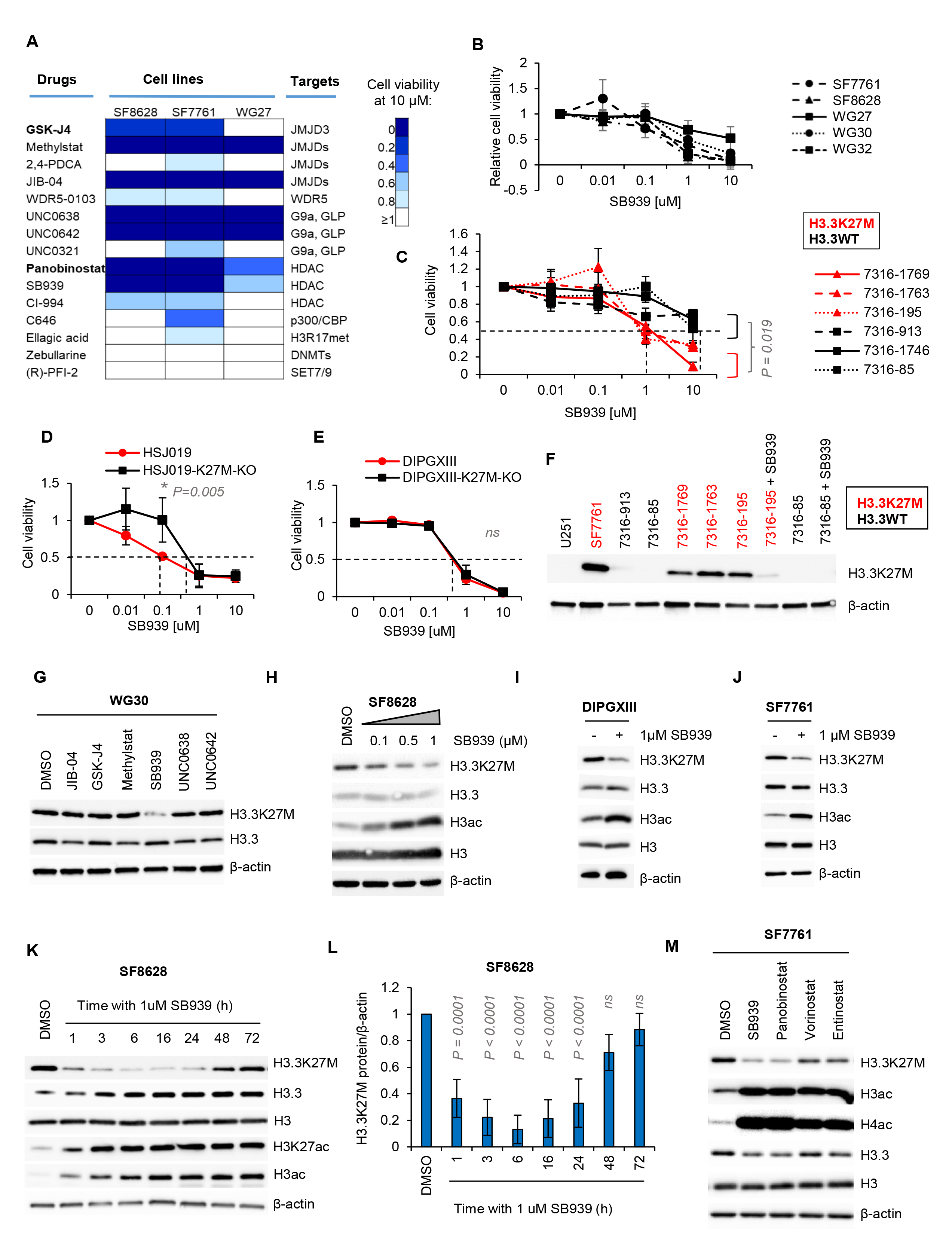
Epigenetic inhibitor screening in H3K27M-expressing paediatric high-grade glioma cells revealed HDACi-dependent decrease of the H3.3K27M oncohistone. **A.** SF8628, SF7761 and WG27 cells were treated for 72 hours with the indicated drugs (10 µM) and subjected to cell viability assay (MTT metabolism). Darker blue colours (legend closer to 0) show stronger cell killing by a particular drug. GSK-J4 and Panobinostat are shown as positive controls. **B.** SF7761, SF8628, WG27, WG30, WG32 cell lines expressing H3.3K27M were treated with serial dilutions of SB939, as indicated, and their cell viability was assessed after 72 hours by MTT assay. Mean cell viability and standard deviation (s.d.) is plotted from three independent experiments. **C.** Patient-derived pHGG spheres expressing either H3.3K27M (listed in red) or wild-type (WT) H3.3 (listed in black) were treated with SB939 at increasing cell concentrations. Cell viability was assessed; mean cell viability and s.d. is plotted from three independent experiments. Two-way ANOVA test showed statistically significant difference between the H3.3K27M and H3.3 WT groups of cells (*P*=0.019). Multiple comparison showed significant differences at 1 and 10 µM treatments (adjusted *P* value of 0.002 and 0.003, respectively). **D-E**. HSJ019 and DIPGXIII pHGG-derived cells expressing H3.3K27M and their isogenic pairs with the deleted *H3F3A* mutated allele (K27M-KO) were treated with SB939 at dilutions shown and assessed for cell viability. Data is shown as mean cell viability and s.d. from three independent experiments. Two-way ANOVA test between parental (H3.3K27M^+^) and K27M-KO cells shows statistically significant difference only for HSJ019 cell line in **D** (adjusted *P* = 0.005). **F.** Presence of the histone H3.3 mutation resulting in pH3.3K27M substitution was verified by Western blotting in cell lines. 24 hour treatment with 1 µM SB939 was included as indicated. **G.** SB939-dependent decrease in H3.3K27M protein levels in WG30 cells treated with selected drugs from **A** as demonstrated by Western blotting. **H.** A dose-dependent decrease of H3.3K27M protein after 48 hours of SB939 treatment shown by Western blotting. A representative experiment of three independent repeats is shown. **I-J.** The H3.3K27M protein levels 16 hours of SB939 treatment in DIPGXIII (**I**) and SF7761 (**J**) cells as shown by Western blotting. A representative experiment of three independent repeats is shown. **K.** Time-dependent decrease of H3.3K27M levels in response to SB939 treatment as shown by Western blotting. A representative experiment of three independent repeats is shown. **L.** Densitometry of H3.3K27M blots relative to β-actin from (**K)** shows mean and s.d. from three independent experiments. Statistically significant changes (*P*<0.0001) were calculated with one-way ANOVA. Dunnett’s multiple comparisons test shows differences for each time-point in relation to DMSO control (adjusted *P* values indicated on the graph). **M.** Modified histone levels in SF7761 spheres treated for 24 h with HDAC inhibitors: SB939 (1 µM), panobinostat (0.05 µM), Vorinostat (1 µM), Entinostat (1 µM) or DMSO control. Total protein extracts were analysed by Western blotting with the antibodies indicated. A representative blot for 3 independent experiments is shown.

Importantly, while we were confirming the status of the H3.3K27M mutation in the cell line panel, we observed that the treatment with SB939 caused a marked decrease in the levels of the mutated histone (**Figure 1F**). The H3.3K27M antibody used was of a very high quality and specificity towards the mutated histone variant (**Supplementary Figure 1L**). In fact, only the treatment with SB939 caused the decrease in H3.3K27M levels and none of the other tested compounds in the initial screen showed a similar effect (**Figure 1G**).

To our knowledge, this is the first finding of the drug able to pharmacologically reduce the levels of the H3.3K27M protein, thus we have investigated the molecular underpinning of this effect. We found that the SB939-mediated decrease of H3.3K27M levels was dose-dependent (**Figure 1H** and **Supplementary Figure 1M)** and consistent in all tested cell lines, including cells grown as adherent monolayers in FBS-containing medium (SF8628, WG30), cells grown as adherent monolayers in the Neurocult medium without FBS (DIPGXIII), as well as cells cultured as spheres (7316-195, 7316-1763 and SF7761) (**Figure 1G-J** and **Supplementary Figure M-N)**. We then tested the kinetics of H3.3K27M decrease in response to SB939 in two different cell lines and found that downregulation of H3.3K27M was significant as early as one hour after the treatment with SB939, with the strongest decrease observed between 6 and 24 hours of treatment (**Figure 1K-L** and **Supplementary Figure 1O-P)**. The levels of H3.3K27M recovered around 48 hours after the treatment, indicating that H3.3K27M decrease was temporary. The effect of SB939 treatment on the upregulation of the histone 3 acetylation was prolonged up to 72 hours, as shown by the total H3 acetylation and H3K27ac levels (**Figure 1K-L** and **Supplementary Figure 1O-P**).

We then examined whether other HDACi could exert a similar effect on H3.3K27M as did SB939. We treated SF7761 spheres for 24 hours with SB939, panobinostat, vorinostat and entinostat and examined the levels of H3.3K27M, as well as histone acetylation. We found a decrease in H3.3K27M levels in response to all the inhibitors tested, although the effect was weaker in vorinostat and entinostat treated cells, which also coincided with a smaller effect of these drugs on upregulation of H3 and H4 acetylation (**Figure 1M**). A previous study demonstrated that shRNAs towards HDAC1 and HDAC2 were efficient as single treatments in impairing the viability of DIPG cells ^22^. We therefore investigated whether silencing of these HDACs would result in a decrease in H3.3K27M levels. We transfected SF8628 cells with siRNA against HDAC1, HDAC2 or both. While the efficient knock down of HDAC1 and HDAC2 was achieved, the levels of H3.3K27M remained unchanged up to 72 hours post transfection **(Supplementary Figure 1Q)**. In addition, the effect of HDAC1/2 knock-down on histone acetylation levels was very moderate, suggesting that pan-HDAC inhibition and a very robust histone hyper-acetylation might be required in order to facilitate the repression of H3.3K27M (as seen for SB939, panobinostat, vorinostat or entinostat treatment) (**Figure 1M** and **Supplementary Figure 1Q)**.

Next, we verified whether the H3.3K27M protein loss after SB939 treatment can globally affect the H3.3K27M occupancy at the chromatin. We performed a ChIP-seq analysis in two cell lines (SF7761 and DIPGXIII) for the H3.3K27M and H3.3 total enrichment at the chromatin after 16 h of treatment with SB939 (the time which corresponds to the most robust reduction of H3.3K27M). As expected, SB939 caused a global loss of the H3.3K27M occupancy at the chromatin. In contrast, the levels of the total H3.3 remained unchanged or even slightly increased after SB939 treatment (**Figure 2A-B**).

**Figure 2.**
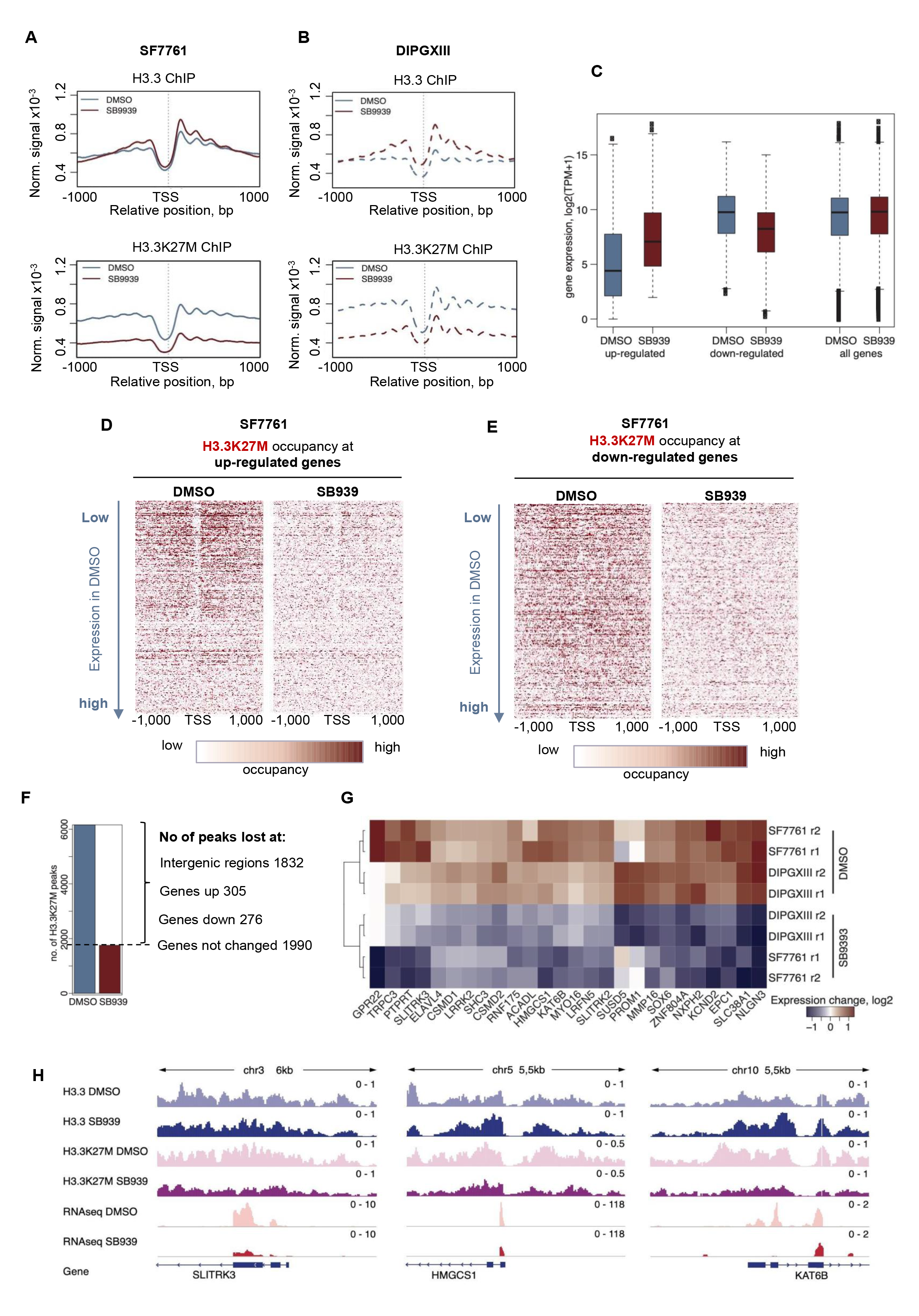
SB939 treatment leads to the loss of H3.3K27M occupancy at the chromatin and specific transcriptomic changes. **A-B**. ChIP-seq results for H3.3 (top panel) and H3.3K37M (bottom panel) profiles around TSS (+/-1 kb) of all protein-coding genes in the SF7761 (**A**) and DIPGXIII (**B**) cells treated with DMSO (blue) or 1 µM SB9393 (red) for 16 hours. **C**. Average gene expression levels in the sets of up- or down-regulated genes in SF7761 and DIPGXIII cells at 16 hours after 1 µM SB939 treatment is shown (red). Gene expression in DMSO controls is shown in blue. **D-E.** Heatmaps showing the H3.3K27M occupancy at up-regulated (**D**) and down-regulated (**E**) genes in SF7761 cells. For each pair, the left and right heatmaps correspond to cells treated for 16 hours with DMSO or 1 µM SB939, respectively. Rows in all heatmaps were ordered by the gene expression level in control DMSO-treated cells (the lowest expression in DMSO at the top). **F.** The number of identified H3.3K27M peaks. The peaks were identified in both cell lines separately and only numbers of overlapped peaks are reported. The blue and the red bars correspond to cells treated for 16 hours with DMSO or SB939, respectively. The genomic distribution is reported for the peaks lost after the SB939 treatment. **G.** Heatmap showing normalised (z-scores) gene expression of 26 genes positively associated with the presence of the H3.3K27M oncohistone based on Bender et al. ^11^. Treatment with SB939 shows decreased expression of these genes (blue). **H.** Example IGV profiles illustrating H3.3K27M loss in SF7761 cells after SB939 treatment at down-regulated genes are shown. Profiles of H3.3, H3.3K27M ChIP-seq and RNA-seq are shown for DMSO or SB939 treatment. Profiles were scaled for each gene locus individually.

The presence of H3K27M mutation was previously shown to drive specific transcriptomic programs, often referred as ‘H3K27M signatures’ ^9, 11^. Therefore we explored gene expression changes induced by SB939 treatment in SF7761 and DIPGXIII cells by RNA sequencing. We hypothesised that the loss of H3K27M in response to SB939 could reverse these ‘signatures’. The PCA analysis showed a clear separation of samples based on the cell lines and SB939 treatment (**Supplementary Figure 2A**). As we saw a good correlation in SB939-triggered gene expression changes between the two cell lines tested (Spearman’s correlation=0.68 and Pearson’s correlation=0.69), we continued further analyses on data integrated from these two cell lines (**Supplementary Figure 2B**). We identified 2,148 upregulated and 1,557 downregulated genes in SB939-treated cells (**Supplementary Figure 2C**). Interestingly, the genes that were up-regulated after SB939 treatment had a very low basal expression in control cells (DMSO-treated) in comparison to the ones that became downregulated or to the average expression of all the genes (**Figure 2C**). This observation suggests that SB939 specifically “unlocked” the genes with the low constitutive expression, while diminishing the expression of highly expressed genes. We analysed the chromatin occupancy of H3.3K27M at the down and up-regulated genes. The loss of H3.3K27M from the chromatin occurred at both up-and down-regulated genes (**Figure 2D-E**). However, for the up-regulated genes the H3.3K27M occupancy was the highest at the ones with the smallest constitutive expression in control cells (**Figure 2D** and **Supplementary Figure 2D)**. This dependency was not seen at the genes downregulated after SB939 treatment (**Figure 2E and Supplementary Figure 2E)**.

To explore functionality of these genes, we performed a Gene Ontology (GO) analysis on SB939 down-and up-regulated genes, and we identified multiple pathways enriched in both groups, respectively (**Supplementary Figure 2F-G**). We have also compared the gene expression induced here with SB939 and in the publicly available RNAseq datasets induced with panobinostat treatment. We found a significant correlation between the gene expression changes induced by SB939 and by panobinostat in DIPGXIII and BT245 cells (Pearson-s correlation=0.41/Spearman’s correlation=0.43 for downregulated genes and Pearson-s correlation=0.55/Spearman’s correlation=0.51 for upregulated genes) (**Supplementary Figure 2H**).

We then investigated the H3.3K27M enrichment at genes that became up or downregulated and which enrichment was lost after SB939 treatment. We detected 6,160 enriched H3.3K27M regions common for the two cells lines in DMSO-treated samples, of which 4,403 disappeared after SB939 treatment. After annotation to the coding genes, we found that these enrichments disappeared at 305 upregulated and 276 downregulated genes in SB939 treated cells (**Figure 2F**). However, considering that more genes were upregulated (2,148) than downregulated (1,557) (**Supplementary Figure 2C)**, the loss of H3K27M enriched sites was more enriched and statistically significant only in downregulated genes. The GO enrichment analysis of these down-regulated genes with the H3.3K27M loss identified numerous pathways, including neuron projection development, plasma membrane bounded cell projection morphogenesis, cell projection morphogenesis, neuron projection morphogenesis, cell morphogenesis involved in differentiation, neurogenesis, generation of neurons, neuron differentiation, axon guidance, cell morphogenesis involved in neuron differentiation, etc (**Supplementary Figure 3A**). Those pathways have been previously linked to the H3K27M oncohistone in gliomas ^11^. Interrogation of our data with the known “H3K27M signatures” encompassing 143 genes previously reported as upregulated in H3.3K27M-expressing pHGGs revealed that out of the 143 H3.3K27M-dependent genes, 26 were down-regulated in SB939- treated samples in both cell lines, including the synaptic adhesion molecule neuroligin-3 (*NLGN3*), which promotes pHGG growth (Fisher’s exact Pv=3e-04) (**Figure 2G-H** and **Supplementary Figure 3B**, the heatmap for all 143 genes is in the **Supplementary Figure 3C**) ^11, 31, 32^. These data show that SB939 treatment, which leads to the downregulation of H3.3K27M, also leads to the loss of a group of genes previously reported as H3K27M-dependent ^11^. However, it is important to stress that the effect of HDAC inhibition on gene expression might be stronger than the effect of the loss of H3.3K27M during HDAC inhibition, which could explain why not all 143 H3.3K27M-dependent genes were regulated in the same way after HDACi treatment (**Figure 2G** and **Supplementary Figure 3C**).

Next, we investigated the mechanism of the H3.3K27M loss in response to SB939. First, we tested whether SB939 caused the transcriptional repression of the *H3F3A* gene, in which one allele contains the mutation encoding the H3.3K27M histone variant. The qPCR analysis showed increased expression of *H3F3A* and *H3F3B* genes in response to SB939 treatment (**Figure 3A-B**). A similar result was observed in RNAseq data from cells treated with SB939 (**Supplementary Figure 4A**) as well as in previously published RNAseq data sets for cell lines treated with panobinostat (**Supplementary Figure 4B-C**). These data suggest that the reduction of H3.3K27M in response to SB939 treatment most likely occurs directly at the protein level and not as a result of decreased mRNA expression. To test this hypothesis, we overexpressed the construct coding for N-terminally Flag-tagged H3.3K27M (Flag-H3.3K27M) in the LN18 glioma cells. As shown by Western blotting with anti-H3.3K27M and anti-Flag antibodies, the signal from the Flag-H3.3K27M band decreased in response to SB939 treatment with both antibodies, while the mRNA expression of the construct remained unchanged upon the treatment (**Figure 3C-D**). To determine whether the wild type H3.3 is similarly affected by SB939 treatment as the mutated histone, we overexpressed the Flag-H3.3K27M and Flag-H3.3WT, in parallel. Only downregulation of the mutated Flag-H3.3K27M was apparent, but not the wild-type Flag-H3.3 or endogenous H3.3 (**Figure 3E**). This result suggests that SB939 treatment specifically reduces the H3.3K27M protein in glioma cells.

**Figure 3.**
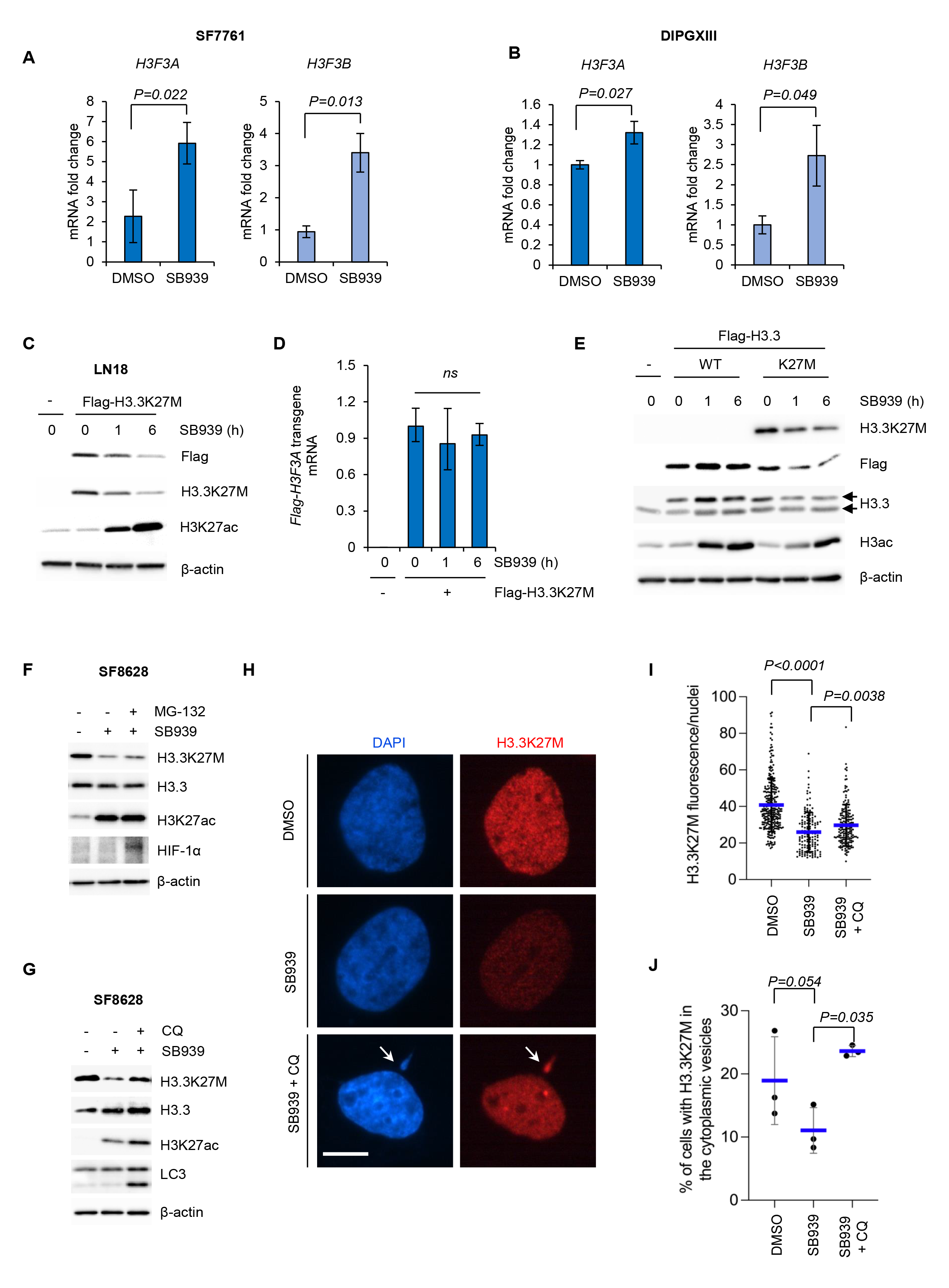
SB939-dependent loss of H3.3K27M is irrespective of *H3F3A* mRNA expression and is partially reversed via chloroquine treatment. **A-B.** Expression of *H3F3A* and *H3F3B* mRNA after 16 hours of treatment with SB939 was tested with qPCR in SF7761 (A) and DIPGXIII (B) cells. *GAPDH* was used as a housekeeping gene. Mean expression and s.d. is shown from three independent experiments. Two-tailed student’s t-tests determined the statistical significance, as indicated with *P* values. **C.** LN18 cells were transfected with Flag-H3.3K27M and 48 hours later exposed to 1 µM SB939 for the times indicated. Western blotting with anti-Flag and anti-H3.3K27M antibodies indicates downregulation of the H3.3K27M protein.. **D.** Expression of Flag-H3.3K27M transgene in experiment shown in C was confirmed by qPCR with forward primer containing Flag sequence and reverse primer binding within *H3F3A* coding sequence. *GAPDH* was used as a housekeeping gene. **E.** LN18 cells were transfected either with Flag-H3.3K27M of Flag-H3.3WT (as indicated) and 48 hours later exposed to 1 µM SB939 for the indicated times. Western blotting with anti-Flag, anti-H3.3K27M And anti-H3.3 antibodies shows expression of transfected histones, accordingly. The top arrow at the H3.3 blot indicates the overexpressed flag-tagged protein and the lower arrow indicates the endogenous H3.3 protein. A representative experiment from 3 independent repeats is shown. **F.** SF8628 cells were treated for 1h with 1 µM SB939 in the presence or absence of 1 µM MG- 132 proteasome inhibitor. Western blotting was performed with the indicated antibodies. HIF- 1α rescue with MG-132 was shown as a positive control for the proteasome inhibition. A representative experiment is shown for three independent repeats. **G.** SF8628 cells were treated for 1h with 1 µM SB939 in the presence or absence of 200 µM chloroquine (CQ). Western blotting was performed with the indicated antibodies. LC3 lipidation is shown as a positive control for CQ treatment. A representative experiment is shown for three independent repeats. **H.** Cells were seeded on coverslips and 24 hours later were exposed to SB939 and CQ treatment for 1 hour, as indicated. Immunofluorescent staining of H3.3K27M (red) and nuclei (DAPI, blue) was performed, as shown with the representative images. White arrows in the bottom panel indicate example vesicles quantified in **J**. Scale bar, 10 µm. **I.** Quantitation of nuclear fluorescence intensity of H3.3K27M staining from **H** was performed for three independent experiments and over 100 of cells were analysed in each condition. Mean intensity per cell (blue line) and standard deviation (black error bars) are overlayed on the top of all measured nuclei in each condition. Unpaired student’s t-test was used to determine statistical significance, as indicated with *P* values. **J.** A graph showing % of cells with extranuclear vesicles positive for DNA (DAPI, blue) and H3.3K27M (red) in each treatment condition (example vesicles indicated with white arrows in **H**). A mean % from three independent experiments (blue line) and standard deviation are shown. Paired student’s t-test was used to determine statistical significance, as indicated with *P* values.

To get more insight, we examined which of two major protein degradation pathways could be responsible for the potential reduction of H3.3K27M levels upon SB939 treatment. The treatment with a proteasomal inhibitor, MG-132, had no significant effect on the levels of the mutated histone in SB939-treated cells (**Figure 4F**). MG-132 stabilised the expression of HIF-1α protein, which is targeted for degradation via E3 ubiquitin ligase VHL and proteasome, confirming that the proteasome degradation pathway has been inhibited (**Figure 3F**). We then tested autophagosome/lysosomal pathway for the potential degradation system of H3.3K27M. Interestingly, the addition of chloroquine (CQ), which impairs lysosomal degradation by increasing the pH of lysosomes, partially restored the levels of H3.3K27M decreased by SB939 (**Figure 3G**).

**Figure 4.**
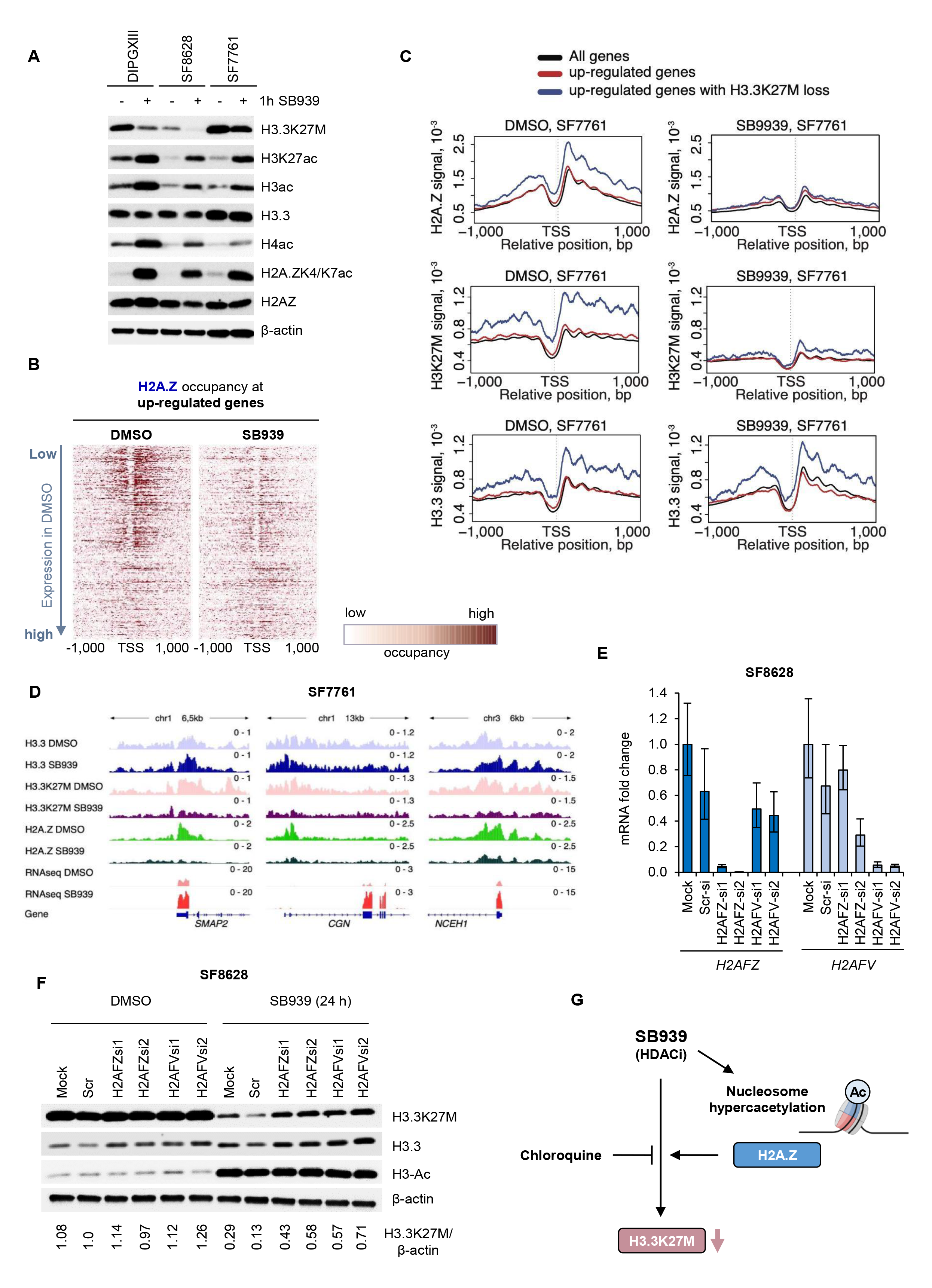
Presence of H2A.Z predisposes pHGG cells to the loss of H3.3K27M during HDAC inhibition with SB939. **A.** DIPGXIII, SF8628 and SF7761 cells were exposed to 1 µM SB939 treatment for one hour and samples were subjected to Western blotting with the indicated antibodies. A representative blots of at least three independent experiments are shown. **B.** Heatmaps showing the H2A.Z occupancy around transcriptional start sites (TSSs) in up-regulated genes in SF7761 cells. The left and right heatmap corresponds to DMSO and SB939 treatment, respectively. Rows in heatmaps were ordered by the gene expression level in control DMSO-treated cells (the lowest expression in DMSO at the top). **C.** The histone variant profiles around TSS in SF7761 cells treated with SB939 for 16 hours. The average H2A.Z (top panel), H3.3K27M (middle panel) and H3.3 (bottom panel) profiles around TSS (+/-2kb) of all protein-coding genes (black line), all up-regulated genes (red), and up-regulated genes with H3.3K27M loss after SB939 treatment (blue) are shown. The left and right plots correspond to control cells and cells after SB939 treatment, respectively. **D.** Examples illustrating H3.3K27M and H2A.Z loss in SF7761 cells after SB939 treatment at up-regulated genes. Profiles of H3.3, H3.3K27M, H2A.Z ChIP-seq and RNA-seq at the loci encompassing selected up-regulated genes. Profiles were scaled for each gene locus individually. **E.** SF8628 cells were double-transfected with siRNAs against *H2AFZ* and *H2AFV* transcripts, control siRNA (Scr-si) or Mock (no siRNA). Efficient knock-down of *H2AFZ* and *H2AFV* at 72 hours post-transfection was verified by qPCR with specific primers in relation to *GAPDH* housekeeping gene. **F.** SF8628 cells were transfected as in **E** and 48 hours post-transfection cells were treated with 1 µM SB939 or DMSO for 24 hours. Samples were then subjected to Western blotting with the indicated antibodies. A representative experiment from three independent repeats is shown. The bottom panel shows quantitation of H3.3K27M densitometry normalised to β-actin. **G.** A scheme showing SB939-dependent downregulation of H3.3K27M in DIPG cells. Upon treatment with SB939, histones present in the nucleosomes undergo rapid hyperacetylation, including the histone variant H2A.Z. Co-occurrence of H2A.Z predisposes cells to the SB939- mediated H3.3K27M loss. The loss of H3.3K27M by SB939 is blocked in the presence of chloroquine, a lysosomal inhibitor and DNA-intercalating agent.

Previous reports have shown that fragments of chromatin can be cleared via lysosomal degradation under certain circumstances, however these studies did not investigate the effect of HDAC inhibition on this process ^33, 34^. We examined the cellular localisation of endogenous H3.3K27M via immunofluorescent staining and asked how it changed in response to SB939 treatment and after the rescue with CQ. The nuclear intensity of H3.3K27M was significantly reduced upon the treatment with SB939 and rescue of the nuclear H3.3K27M was visible when CQ was present during the SB939 treatment (**Figure 3H-I**). In addition, we observed that in each condition, a fraction of cells had extracellular vesicles present in the cytoplasm that contained DNA (stained with DAPI) and H3.3K27M (as indicated in **Figure 3H** with arrows in the representative picture for SB939+CQ). We determined the percentage of cells in each condition with visible cytoplasmic vesicles. While cells with these vesicles were the least abundant in the SB939-treated group, these were significantly increased when the CQ was present (**Figure 3J**). This result suggests that CQ impairs the clearance of the chromatin (and H3.3K27M) in these vesicles exported extranuclearly. However, the nuclear rescue of H3.3K27M was also visible in SB939+CQ treated samples (**Figure 3I**), implying that some CQ-dependent direct or indirect mechanisms of H3.3K27M downregulation in the nuclei also existed upon SB939 treatment.

While our data supported the notion that chromatin fragments containing H3.3K27M were at least partially cleared via lysosomes, it was difficult to explain why this process was so rapid and temporary, and whether this would be the only clearance pathway of the mutated histone. Therefore, we investigated more thoroughly the events coinciding with the H3.3K27 loss triggered by SB939 treatment. As expected for the HDACi, the treatment with SB939 caused an immediate and significant increase in the acetylation of numerous histones, including H3, H4, but also H2A.Z histone variant (**Figure 4A**). Such increased histone acetylation could potentially cause destabilisation of nucleosomes and chromatin relaxation ^35^. Previous studies showed that incorporation of the H2A.Z and H3.3 histone variants in the same nucleosomes made them more unstable than the presence of canonical H2A and H3 histone variants, although the significance of H2A.Z histone variant in the nucleosome stability in H3.3K27M-expressing DIPG cells has not been investigated to date ^36, 37^. As our antibody against H2A.Z does not distinguish between H2A.Z.1 (encoded by *H2AFZ*) and H2A.Z.2 (encoded by *H2AFV*) isoforms, we have investigated the expression of both these genes upon SB939 treatment in various cell lines using qPCR or in RNAseq datasets from SB939 or panobinostat treated cells. We have found that across all analysed datasets expression of the more abundant isoform, *H2AFZ*, was decreased in response to HDACi in most datasets tested, while *H2AFV* did not significantly change or was even moderately increased in some cells (**Supplementary Figure 5A-D**). This result could potentially explain the relatively stable overall levels of the H2A.Z protein detected with the pan-H2A.Z antibody after the SB939 treatment (**Figure 5A**).

We assessed the H2A.Z occupancy at the chromatin and in the context of gene expression by performing the H2A.Z ChIPseq in SF7761 and DIPGXIII cells treated with SB939 or with DMSO. The ChIPseq conditions for H2A.Z were matching the ones showed earlier for H3.3 and H3.3K27M in **Figure 2**. In one of the cell lines (SF7761) SB939 treatment caused average H2A.Z loss in the TSS proximal regions in all the genes, but not in DIPGXIII cells (**Supplementary Figure 6A**). However, a more striking difference in the H2A.Z occupancy in both cell lines was apparent, when up-and down-regulated genes upon SB939 treatment were analysed separately. The highest H2A.Z occupancy was observed at the genes with the lowest expression in DMSO samples and that then became up-regulated by SB939 (**Figure 4B** and **Supplementary Figure 6B**). Moreover, this was more apparent, when we compared H2A.Z occupancy at all genes (black line), up-regulated genes (red line) and up-regulated genes with H3.3K27M loss (blue line) (**Figure 4C/**top row and **Supplementary Figure 6C/**top row). The genes with the highest loss of H2A.Z occupancy where the ones that also lost the H3.3K27M occupancy and became up-regulated by SB939 treatment (**Figure 4C/**middle row panel and **Supplementary Figure 6C** top/middle rows; bottom row with H3.3 WT is shown for the comparison).

The presented data suggest that the co-occurrence of both H3.3K27M and H2A.Z histone variants in close proximity at actively transcribed genes could facilitate their loss, particularly in the situation of increased histone acetylation during HDAC inhibition. Perhaps the presence of H2A.Z promotes the total loss of H3.3K27M in DIPG cells as was shown with Western blotting. To verify this hypothesis, we depleted H2A.Z levels in cells with siRNA and investigated the loss of H3.3K27M after SB939 treatment. SF8628 cells were double transfected with two different siRNAs targeting either *H2AFZ* or *H2AFV* expression to ensure a prolonged knock-down of H2A.Z. Transfected cells were treated with SB939 for 24 hours and we performed qPCR and western blotting analyses. The specificity and effectiveness of silencing was confirmed with qPCR primers detecting both H2A.Z isoforms (**Figure 4E**). Importantly, the knock-down of either *H2AFZ* or *H2AFV* genes consistently impaired the reduction of total H3.3K27M levels upon the SB939 treatment as shown by Western blotting (**Figure 4F**). Similar results were obtained with another DMG cell line (WG32) tested (**Supplementary Figure 6E-F**). Altogether, the presented data support the conclusion that increased histone acetylation (e.g. during HDAC inhibition) promotes the loss of H3.3K27M, particularly in the presence of H2A.Z histone variants (**Figure 4G**).

## DISCUSSION

DIPG childhood gliomas expressing the H3.3K27M oncohistone have the worst prognosis with no curative therapy available. In this study, we demonstrate for the first time that pan-HDAC inhibition leads to a significant decrease in the H3.3K27M protein levels in a number of H3.3K27M^+^ DIPG-derived cells. SB939-dependent decrease of H3.3K27M is very rapid and accompanied by an immediate increase in histone acetylation. We show that co-occurrence of H2A.Z and H3.3K27M at the same genomic locations pre-disposes cells to the HDACi-dependent loss of H3.3K27M, particularly at the genes which will become up-regulated upon SB939 treatment. Depletion of H2A.Z isoforms from H3.3K27M^+^ DIPG cells significantly impairs the SB939-dependent loss of H3.3K27M, emphasising the role of the nucleosome components such as the H2A.Z histone variant in the stability of the H3.3K27M protein. We also show that SB939-dependent loss of H3.3K27M is partially blocked with chloroquine, acting most likely through the inhibition of lysosomal degradation pathway.

Intriguingly, HDAC inhibition with SB939 leads to a preferable loss of the H3.3K27M oncohistone, but not the wild type H3.3 histone variant, which we showed at the total protein levels with Western blotting, but also locally at the genes with ChIPseq analysis. This reduction of the histone H3.3K27M level is not due to decrease of transcription. We demonstrate that both *H3F3A* (one allele of the *H3F3A* gene is wild type) and *H3F3B* mRNA are elevated in response to HDAC inhibition and the total H3.3 protein levels are not changed or increased. The previously reported mass spectrometry analysis showed that mutated H3.3K27M constitutes only between 3 to 17% of the total H3 protein levels, which could explain why the total H3.3 protein remain at similar levels, especially if the *H3F3A* and *H3F3B* mRNA remain constant or are slightly upregulated after the SB939 treatment ^10^. The present study shows that that SB939 triggers cellular mechanisms leading to the loss of H3.3K27M specifically, while sparing the remaining wild type H3 histone pool.

We present two potential mechanisms responsible for selective downregulation of H3.3K27M protein and reduced occupancy at the chromatin. Out of two tested inhibitors of protein degradation pathways, chloroquine partially blocked the loss of H3.3K27M induced by SB939 treatment. It is not entirely clear how chloroquine impairs the SB939-mediated loss of H3.3K27M protein levels as this compound has multiple actions besides being the lysosomal/autophagosome inhibitor. Blockade of the lysosomal turnover of chromatin fragments during SB939 treatments by chloroquine is a plausible explanation for the H3.3K27M loss. This is in line with previous studies showing that H3 can be turned over via lysosomes in the autophagy process, however to date this has not been investigated in the context of HDAC inhibition ^33, 34^. In addition, the lysosomal degradation of H3.3K27M can offer only a partial mechanism of the H3.3K27M loss during HDACi treatment, as our data equally suggests that chloroquine already restores the H3.3K27M levels at the nuclear level. In addition to inhibiting lysosomal degradation, chloroquine has been historically known for its DNA-intercalating properties, which may lead to inhibition of DNA replication and RNA synthesis ^38^. One could speculate that chromatin is stabilised by CQ intercalation and disassembly of nucleosomes impaired, resulting in a weaker H3.3K27M degradation. Nevertheless, this mechanism needs a thorough future investigation.

Histone variants H2A.Z and H3.3, are frequently placed in the same nucleosomes and cooperate to regulate transcription ^39^. As H2A.Z can be an oncogenic driver, we explored the role of H2A.Z in DIPG cells after HDAC inhibition ^40, 41^. We show that many of H2A.Z and H3.3K27M ChIP-seq peaks overlap with each other in DIPG cells at genes with a low constitutive expression. Additionally, both peaks are lost after HDAC inhibition at upregulated genes. The function of H2A.Z in H3.3K27M^+^ DMG tumours and in the context of sharing the nucleosomes with H3.3K27M has not been investigated yet. The unstable H3.3-H2A.Z nucleosomes might be even more affected when H3.3 is substituted with the H3.3K27M oncohistone. However, it is equally possible that the presence of H2A.Z, and particularly its hyperacetylated form, might recruit some additional proteins, like RNA polymerase II and other partners that will additionally alter gene expression or induce turnover of H3.3K27M containing nucleosomes ^40^. Future studies will focus on precise dissecting of the H2A.Z role in the H3.3K27M stability and turnover.

In summary, this is a first demonstration that decreasing the levels of the detrimental H3.3K27M oncohistone can be achieved pharmacologically with HDAC inhibitors. While this discovery does not offer an immediate therapeutic solution to improve the outcome of DMG patients, these results open some alternative pathways to target H3.3K27M. The ultimate future goal is to explore the potential of such mechanisms in H3.3K27M-driven tumorigenesis.

## MATERIALS AND METHODS

### Cell lines and reagents

A summary of pHGG cell lines used in this study is shown in Supplementary Table 1. SF8628 and SF7761 cells were a kind gift from Dr Rintaro Hashizume and were cultured as described ^28, 42, 43^. BT245, SU-DIPG-XIII and HSJ019 were a kind gift from Dr Nada Jabado (McGill University) and were cultured as adherent cells on poly-L-Ornithine and Laminin coated flasks in medium containing Neurocult NS-A Basal Medium and Supplement, rhEGF (20 ng/ml), bFGF (10 ng/ml) and Heparin (2.5 µg/ml). ^19, 29, 30^. 7316-1769, 7316-1763, 7316-195, 7316-913, 7316-1746 and 7316-85 pHGG cells were obtained from the Children’s Brain Tumor Tissue Consortium (CBTTC, via The Children’s Hospital of Philadelphia (CHOP)) and were cultured as spheres in DMEM/F12, GlutaMAX, B27 Supplement w/o Vitamin A, N2 supplement, rhEGF (20 ng/ml), bFGF (20 ng/ml) and 2.5 µg/ml Heparin. All media were supplemented with penicillin and streptomycin antibiotics. WG27, WG30 and WG32 cells were derived from pHGG patient biopsies collected at the Children’s Memorial Health Institute (CMHI) in Warsaw as described previously ^44^. The use of tumour biopsies was approved by the Research Ethics Committee at CMHI in Warsaw, Poland, and informed consents were obtained from the patients. Biopsies were placed in DMEM/F12 medium and transported on ice from the hospital. Tissues were dissected mechanically with a scalpel on a Petri dish in 0.5 ml of DMEM/F12 medium until homogenous tissue suspension was achieved. After filtering through 100- and 40-micron cell strainers cells were centrifuged and washed with fresh DMEM/F-12. Cells were resuspended in DMEM/F-12 medium supplemented with 10% foetal bovine serum and anti-mycotic antibiotics and cultured as adherent cells. Cell proliferation was monitored. Expression of the H3.3K27M mutation in obtained well-proliferating cultures was determined with Western blotting. Drugs used in epigenetic screen in the Figure 1A were from the Epigenetic Screening Library (11076, Cayman). Chloroquine (C6628) and MG-132 (474787) were from Sigma-Aldrich.

### Plasmid and siRNA transfection

The plasmid expressing flag-H3.3 (pcDNA4/TO-Flag-H3.3) was purchased from Addgene (47980). The plasmid expressing Flag-H3.3K27M was generated by site-directed mutagenesis of flag-H3.3 plasmid using primers: F: 5’-GAGTGCGCCCTCTACTGG-3’ and R: 5’- ATGCGAGCGGCTTTTGTAG-3’. Successful mutagenesis was confirmed by sequencing at Genomed S.A. (Warsaw, Poland). Plasmid transfections into LN18 cells were performed with jetPRIME transfection reagent (114-07, PolyPlus), according to manufacturer’s instructions. siRNA transfections in adherent SF8628 and WG32 cells were performed with DharmaFECT 1 transfection reagent (Dharmacon) according to manufacturer’s protocol using the siRNA duplexes listed in the Supplementary Table 2.

### Western blotting

Cells were washed in PBS and lysed in SDS lysis buffer (10 mM Tris-Cl, pH 7.5, 0.1 mM EDTA, 0.1 mM EGTA, 0.5% SDS, 0.1 mM β-mercaptoethanol, protease/phosphatase inhibitors), followed by sonication and centrifugation. Lysates with equal amount of proteins were subjected to SDS-PAGE and western blotting. The primary antibodies included: H3 total (ab1791, Abcam), H3.3 (09-838, Millipore), H3.3K27M (61803, Active Motif), H2A.Z (ab150402, Abcam), H2A.ZK4/K7ac (#75336, Cell Signalling Technologies (CST)), H3ac (06-599, Millipore), H3K27ac (8173S, CST), H4ac (06-866, Millipore), cleaved caspase 3 (#9661S, CST), cleaved caspase 7 (#9491S, CST), PARP (9542, CST), HDAC1 (09-720, Millipore), HDAC2 (05-814, Millipore), Flag (F3165, Sigma-Aldrich), LC3 (L7543, Sigma-Aldrich), HIF-1α (ab179483, Abcam), β-actin (A3854, Sigma-Aldrich). An enhanced chemiluminescence detection system (ECL) and Chemidoc (BioRad) were used to develop the signal from HRP-conjugated secondary antibodies.

### qPCR

Glioma cells were treated accordingly and total RNA was isolated with RNeasy Plus Kit (Qiagen) according to manufacturer’s instructions; RNA concentrations and purity were assessed with NanoDrop 2000 (Thermo Fisher Scientific). Equal amounts of total RNA per each condition were used to make cDNA with SuperScript™ III Reverse Transcriptase kit (Invitrogen) and oligo-dT primers. PCR reactions with technical triplicates were performed with 2× Fast SYBR GREEN PCR Master Mix (Applied Biosystems) using QuantStudio 12K Flex equipment (Applied Biosystems, Life Technologies). The qPCR primer sequences are listed in the Supplementary Table 3.

### Cell viability and cell proliferation assays

For cell viability, MTT assay was performed. 5×10^3^ cells were seeded onto 96-well plates and cells that were cultured as spheres were supplemented with 5% FBS to attach for the time of the assay, as previously indicated ^22^. Next day, the drug dilutions were added in technical triplicates. After 72 hours, the MTT solution (0.5 mg/mL; Sigma-Aldrich) was added and after 1 hour of incubation at 37 °C was replaced with 100 µl of DMSO to dissolve water-insoluble dark blue formazan crystals present in the attached cells. Optical densities were measured at 570 nm and 620 nm using a scanning multi-well spectrophotometer. For cell proliferation, cells were seeded at density of 5×10^3^ per well in 96-well plates. Cells were treated with dilutions of SB939 for 24 or 48 hours and cell proliferation was assessed using ELISA BrdU kit (Roche Diagnostics GmbH), according to manufacturer’s protocol.

### Immunofluorescence

SF8628 cells were seeded on glass coverslips in 24-well plates at density of 2×10^4^ cells per well, and the next day 1 hour treatment with SB939 in the presence or absence of chloroquine was performed. Cells were washed with PBS, fixed for 10 minutes with 4% PFA, followed by a triple wash with PBS. This was followed by 10-minute permeabilization with ice-cold 100% methanol and wash in PBST. After 1 hour blocking (3% donkey serum, 1% BSA, 0.3% Triton X-100, PBS) the H3.3K27M antibody (1:500, 61803, Active Motif) diluted in a blocking solution was added and incubated for 1 hour. This was followed by a triple wash in PBST and an hour incubation with a secondary donkey anti-rabbit antibody conjugated with Alexa Fluor-555 (1:2000, A31572, Invitrogen). Coverslips were washed 3x with PBST and once with distilled water, followed by mounting with Vectashield mounting solution with DAPI (Vector Laboratories).

### RNAseq

RNA was isolated using RNeasy Plus Kit (Qiagen) according to manufacturer’s instructions and the quality was assessed with the Agilent 2100 Bioanalyzer with an RNA 6000 Pico Kit (Agilent Technologies). PolyA-enriched RNA libraries were prepared with the KAPA Stranded mRNA Sample Preparation Kit (Kapa Biosystems).

Sequencing was performed with NovaSeq 6000 (Illumina) at a depth of 18-50 million paired reads per sample. The sequenced paired-end reads were mapped to hg38 genome using tophat2 aligner v2.1.1 with the default parameters. The expression estimates for each gene were obtained using Bioconductor package DESeq2. TPM (Transcripts per Kilobase Million) values were calculated and used to perform all visualizations and all analyses besides differential expression. Genes that had significant (Benjamini and Hochberg-corrected P <0.05 and |log2(Fold Change)|>1) changes in their expression levels were called as differentially expressed. All gene ontology analyses were performed with GO.db Bioconductor package and 0.05 significance threshold on Bonferroni corrected P-value (Fisher’s exact test). All statistical analyses were performed in R programming environment (http://r-project.org). The H3.3K27M signature was obtained from Bender et al. ^11^. External data sets were downloaded from the GEO database for cells treated with panobinostat. We used expression profiles for BT245 and DIPGXIII from GSE117446 ^30^, and DIPGIV and DIPGXIII from GSE94259 ^25^. The RNAseq data generated in this project were deposited at the NCBI platform under GSE232283 Number.

### Chromatin immunoprecipitation sequencing (ChIPseq)

ChIP method was adapted from Cook *et al.,* with some modifications ^45^. DIPGXIII or SF7761 cells were treated with DMSO or SB939 for 16 hours and collected for fixation with 1% formaldehyde and neutralisation with 0.125 M glycine. Cell pellets were washed twice with ice-cold PBS supplemented with protease inhibitor cocktail. 1.5×10^6^ cells were used per each IP. Cells were resuspended in 15 ml of ice-cold buffer A (0.32 M sucrose, 15 mM HEPES, pH 7.9, 60 mM KCl, 2 mM EDTA, 0.5 mM EGTA, 0.5% (w/v) BSA, 0.5 mM DTT, protease inhibitors cocktail) and incubated for 20 min on ice. Subsequently, nuclei were released by passing cell suspension through the Dounce homogenizer (20 strokes) and layering it over 15 ml of buffer A+ (1.3 M sucrose, 15 mM HEPES, pH 7.9, 60 mM KCl, 2 mM EDTA, 0.5 mM EGTA, 0.5% (w/v) BSA, 0.5 mM DTT, protease inhibitors cocktail) in 50 ml falcon tube. After 5 min of centrifugation at 1000 x g and 4 °C, nuclear pellets were washed twice with 10 ml of buffer W (10 mM Tris·Cl, pH 7.4, 15 mM NaCl, 60 mM KCl, protease inhibitors cocktail) and pelleted by 350 x g centrifugation for 4 min at 4 °C. Nuclei were resuspended with 400 µl of buffer W and supplemented with final 1.2 mM CaCl_2_. Subsequently, 100 µl of nuclease-free water containing 20 units of micrococcal nuclease (MNase, Worthington) was added and enzymatic chromatin digestion was carried out for 25 min at 25°C with 1000 rpm shaking. Reaction was then stopped with 30 µl of 0.5M EDTA and EGTA solutions. Subsequently, chromatin was solubilised with supplementation of 0.5% SDS followed by 10-fold dilution in LB3 buffer (1 mM EDTA, 10 mM Tris·Cl, pH 7.5, 1% (w/v) sodium deoxycholate, 0.5% (w/v) sarkosyl, 1% (v/v) Triton X-100). After 10 min incubation on ice, samples were centrifuged for 10 min at 16000 x g 4 °C and supernatant collected. 200 µl of lysate was taken for input sample and the rest used in over-night immunoprecipitation with 4 µg of antibody and end-over-end rotation at 4 °C. The following antibodies were used for each IP reaction: H3.3 (09- 838, Millipore), H3.3K27M (61803, Active Motif) and H2A.Z (ab150402, Abcam). The next day 20 µl of pre-washed Protein A Dynabeads (Invitrogen) were added, and incubation continued for another 2 hours at 4 °C. Beads were then washed 6 times with LB3 buffer and collected using the magnetic stand. Immunoprecipitated chromatin was eluted twice with 125 µl of elution buffer (0.2% (w/v) SDS, 0.1 M NaHCO3, 5 mM DTT) at 65 °C for 10 min. Next, chromatin eluates and inputs were subjected first to reverse crosslinking over night at 65 °C, then to 30 min of RNAse A treatment at 37°C followed by Proteinase K treatment for 4 hours at 50 °C. DNA was then extracted with phenol/chloroform/isoamyl alcohol reagent as previously described and precipitated from the aqueous phase with 3 volumes of 100% ethanol in the presence of 0.1 volume of 3M sodium acetate and 1 ul of glycogen ^46^. DNA was additionally purified with ZymoResearch DNA Clean & Concentrator-5 columns (D4003T) and digested DNA profiles confirmed using the 2100 Bioanalyzer with the Agilent DNA High Sensitivity chip (Agilent Technologies). The libraries were then prepared using the Qiaseq Ultra Low Input Library Kit (Qiagen), according to manufacturer’s instructions. Sequencing was performed with NovaSeq 6000 or HiSeq1500 (Illumina) at a depth of 20 to 56 million single-end reads.

The paired-end fragments were generated. Reads were mapped to the to hg38 genome using Bowtie aligner version 0.12.9. Only uniquely mapped reads were retained. The reads with insert sizes <50 bp or >500 bp were removed from further analysis. Genomic positions with the numbers of mapped tags above the significance threshold of Z-score of 7 were identified as anomalous, and the tags mapped to such positions were discarded ^47^. H2A.Z, H3.3 and H3.3K27M occupancies were estimated as tag frequencies. Input correction was not used in line with previous study ^48^. Enrichment peaks were called with MACS2.0 (q-value< 0.0001 and log2(Fold Change > 2)). The ChIPseq data generated in this project was deposited at NCBI under GSE232283 Number.

### Statistical analysis of biochemical data

Statistical analyses for qPCRs, immunofluorescent intensity, MTT, BrdU and western blotting quantitation were performed with Student’s t-test and/or ANOVA using GraphPad Prism software (GraphPad Software Inc.), as indicated in figure legends.

### COMPETING INTERESTS

The authors declare no competing interests.

### AUTHORS CONTRIBUTIONS

KBL conceived, designed, performed, supervised experiments and interpreted the results; and wrote the manuscript. JM conceived the project and secured the funding; conceived, designed or supervised the experiments; performed computational analyses and interpreted the results; wrote the manuscript. APDF, CJ, MD, KW, BG, PS and BK (Kaza) performed the experiments. MKC, WG and JT provided pHGG biopsies from patients to derive cell lines and performed the molecular analysis of biopsies. BK (Kaminska) provided guidance on the project, infrastructure for the experiments and co-wrote the manuscript.

## ACKNOWLEDGEMENTS

Thank you to Prof. Ester Hammond (University of Oxford) for the critical reading of the manuscript. We acknowledge Sequencing Lab at the Nencki Institute for sequencing all NGS samples.

## FUNDING

This work was supported by the National Science Center, Poland grant (award no. 2017/27/B/NZ2/02827) to JM. Part of JM’s and KBL’s salaries were founded by Foundation for Polish Science (FNP) under the International Research Agendas Program (grant number MAB/2018/6) and the National Science Center, Poland grant (award no. 2019/33/B/NZ1/01556), respectively.

## Notes

### Competing Interest Statement

The authors have declared no competing interest.

