## Supplementary Figures and Methods for "H2A.Z histone variants facilitate HDACi-dependent removal of H3.3K27M mutant protein in paediatric high-grade glioma cells"

Table of contents: Page

Supplementary Figure Legends 2

Supplementary Tables 8

Supplementary Table 1. Cell lines 8

Supplementary Table 2. siRNA sequences 9

Supplementary Table 3. RT-PCR primers 9

### Supplementary Figure Legends

**Supplementary Figure 1. Characterisation of cellular response to inhibition of epigenetic modifiers.**

**A-E.** SF7761, SF8628, WG27, WG30 and WG32 were treated with 10-fold dilutions of the following drugs: GSK-J4 (**A**), JIB-04 (**B**), Methylstat (**C**), UNC0638 (**D**) and UNC0642 (**E**) for 72 hours and relative cell viability was assessed with MTT assay. Mean cell viability and standard deviation is plotted in relation to DMSO control from 3 independent biological repeats.

**F-G.** Cell lysates of SF8621, WG27 (F) and WG32, WG30, SF8628 (G) cells were subjected to western blotting and expression of H3.3K27M was determined, as shown.

**H.** SF8628 cells were treated with 1 μM SB939 for the times indicated and cellular lysates were subjected to western blotting to detect apoptosis markers, such as cleaved (cl.) PARP, Caspase-3 and Caspase-7. A representative experiment from three independent repeats is shown.

**I.** SF8628 cells were exposed to dilutions of SB939 for 24 or 48 hours and cell proliferation was assessed with BrdU assay. Mean cell proliferation (normalised to DMSO control) is shown with standard deviation from three independent experiments.

**J.** BT245 pHGG-derived cells expressing H3.3K27M and their isogenic pair with deleted *H3F3A* mutated allele (K27M-KO) were subjected to the treatment with SB939 at dilutions shown and assessed for cell viability. Mean cell viability and standard deviation is plotted from three independent experiments. No statistically significant (ns) difference in cell viability was observed between the two cell lines. Dotted line indicates 50% drop in cell viability in relation to DMSO control.

**K.** Normal human astrocytes (NHA) were treated with 10-fold dilutions of SB939 and 72 hours later assessed for cell viability with MTT assay. Dotted line indicates 50% drop in cell viability in relation to DMSO control.

**L.** Adult glioblastoma (GBM) wild type for H3 and DIPG positive for H3.3K27M cell extracts were subjected to western blotting and probed with H3.3 (wild-type/total) and H3.3K27M-specific antibodies.

**M.** WG30 cells were exposed to SB939 as indicated for 48 hours and western blotting was performed with the antibodies shown. A representative experiment of three independent repeats is shown.

**N.** 7316-195 and 7316-1763 spheres derived from pHGG samples were incubated with 1 μM SB939 for 16 hours and cell lysates subjected to western blotting with the antibodies shown. A representative experiment of three independent repeats is shown.

**O.** SF7761 spheres were subjected to a time-course treatment with 1 μM SB939 and H3.3K27M levels were determined with the western blotting in relation to other control antibodies used. A representative experiment of three independent repeats is shown.

**P.** Densitometry of H3.3K27M signal from (O) normalised to β-actin is shown as a mean expression from three independent experiments in relation to DMSO treatment.

**Q.** SF8628 cells were transfected with siRNAs specific to HDAC1, HDAC2 or with control non-targeting siRNA (Scr) and 72 hours later were subjected to western blotting analysis with the antibodies, as indicated. A representative experiment of three independent repeats is shown.

**Supplementary Figure 2. RNAseq and ChIPseq analysis of DIPG cell lines treated with SB939**

**A.** Principal Component Analysis demonstrating gene expression differences between treatments and cell lines, as well as similarities of biological replicates.

**B.** Correlation of gene expression ratios (SB939/DMSO) obtained for the SF7761 cell line (x-axis) and DIPGXIII cell line (y-axis). Spearman’s (s.cor) and Pearson’s (p.cor) correlation coefficients are indicated above the plot.

**C.** Volcano plot showing differences in gene expression levels between DMSO control and SB939-treated cells. The red dots correspond to statistically significantly differentially expressed genes (numbers of genes are indicated above the plot).

**D-E.** Heatmaps showing H3.3K27M levels around up-regulated (**D**) and down-regulated (**E**) genes in DIPGXIII cell line. For each pair, the left and the right heatmap corresponds to DMSO and SB939 treatment respectively. Rows in all heatmaps were ordered by the gene expression level in control cells (DMSO).

**F-G.** Top 10 Gene Ontology (GO) terms enriched in the sets of down-regulated (**F**) and up-regulated genes (**G**). The bars correspond to scaled (-log10) p-values. The red lines indicate Bonferroni corrected threshold for significance.

**H.** Scatter plots showing similarities of gene expression changes after SB939- and Panobinostat (PANO) treatments. X- and Y-axes correspond to SB939 and Panobinostat respectively. The left plot shows results obtained for down-regulated genes, while the right plot corresponds to up-regulated genes. The gene expression changes are shown as differences of log-scaled expression estimations.

**Supplementary Figure 3. Characterisation of SB939-dependent genes**

**A.** Top 10 Gene Ontology (GO) terms enriched in the sets of down-regulated genes with H3.3K27M loss after SB939 treatment. The bars correspond to scaled (-log10) p-values. The red lines indicate Bonferroni corrected threshold for significance.

**B.** Examples illustrating H3.3K27M loss in DIPGXIII cells after SB939 treatment at down-regulated genes. Profiles of H3.3, H3.3K27M ChIP-seq and RNA-seq are shown for DMSO or SB939 treatment and were scaled for each gene locus individually.

**C.** Heatmap showing normalized (z-scores) gene expression of entire H3.3K27M signature (143 genes) obtained from Bender et al. 2013 in the RNAseq data in SF7761 and DIPGXIII cells generated in this study.

**Supplementry Figure 4. Expression if *H3F3A* and *H3F3B* in pHGG cells treated with HDACi**

**A.** Expression estimations of *H3F3A* and *H3F3B* genes in RNAseq data obtained for DMSO and SB939-treated SF7761 and DIPGXIII cells, as indicated (16 hour treatments). Blue and red dots represent estimations in individual biological replicates for the control and SB939-treated cells, respectively.

**B.** Expression estimations of *H3F3A* and *H3F3B* genes downloaded from publicly available RNAseq data (GSE117446) in DMSO and Panobinostat (Pano)-treated cells. Top plots correspond to BT245 cell line, while bottom plots correspond to DIPGXIII cell line. Blue and red dots represent estimations in individual biological replicates for the control and Panobinostat-treated cells.

**C.** Expression estimations of *H3F3A* and *H3F3B* genes downloaded from publicly available RNAseq data (GSE94259) in DMSO and Pano-treated cells. Top plots correspond to DIPGXIII cell line, while bottom plots correspond to DIPGXIII cell line. Blue and red dots represent estimations in individual biological replicates for the control and Pano-treated cells.

**Supplementry Figure 5. Expression if *H2AFZ* and *H2AFV* in pHGG cells treated with**

**HDACi**

**A.** Expression of *H2AFZ* and *H2AFV* in DIPGXIII cells treated with SB939 for 16 hours measured by qPCR. *GAPDH* was used as a reference gene.

**B.** Expression of *H2AFZ* and *H2AFV* in SF8628 cells treated with SB939 for 24 hours measured by qPCR. *GAPDH* was used as a reference gene.

**C.** Expression estimations of *H2AFZ* and *H2AFV* genes from RNAseq data obtained for control (DMSO) and 16 hour SB939 treatment of SF7761 or DIPGXIII cells. Blue and red dots represent estimations in individual biological replicates for the control and SB939-treated cells.

**D.** Expression estimations of *H2AFZ* and *H2AFV* genes downloaded from publicly available data (GSE117446) in control (DMSO) and Panobinostat (Pano)-treated BT245 or DIPGXIII cells. Blue and red dots represent estimations in individual biological replicates for the control and Pano-treated cells.

**E.** Expression estimations of *H2AFZ* and *H2AFV* genes downloaded from publicly available data (GSE94259) in control (DMSO) and Pano-treated DIPGVI or DIPGXIII cells. Blue and red dots represent estimations in individual biological replicates for the control and Pano-treated cells.

**Supplementry Figure 6. Presence of H2A.Z predisposes pHGG cells to the loss of H3.3K27M during HDAC inhibition with SB939.**

**A.** H2A.Z ChIPseq analysis showing average H2A.Z profiles around TSS sites (+/- 1 kb) of all protein coding genes in SF7761 (line) or DIPGXIII (dash line) cells treated with DMSO (control, blue) or SB939 (red).

**B.** Heatmaps showing H2A.Z occupancy in DIPGXIII cells around TSS sites of SB939-up-regulated genes. The rows were sorted according to gene expression levels in control (DMSO) cells.

**C.** The selected histone variant profiles around transcriptional start sites (TSSs). The average H2A.Z (top), H3.3K27M (middle) and H3.3 (top) (bottom) profiles around TSS (+/- 1 kb) of all protein-coding genes (black line), all up-regulated genes (red), and up-regulated genes with H3.3K27M loss after SB939 treatment (blue). The left and right plots correspond to control cells and cells after SB939 treatment respectively. The profiles are obtained for DIPGXIII cells.

**D.** Examples illustrating H3.3K27M and H2A.Z loss in SF7761 cells after SB939 treatment at up-regulated genes. Profiles of H3.3, H3.3K27M, H2A.Z and RNA-seq at the loci encompassing selected SB939-up-regulated genes in DIPGXIII cells. Profiles were scaled for each gene locus individually.

**E.** WG32 cells were double-transfected with siRNAs against *H2AFZ* and *H2AFV* transcripts, control siRNA (Scr-si) or Mock (no siRNA) and efficient knock-down of *H2AFZ* and *H2AFV* at 72 hours post-transfection was verified by qPCR with specific primers in relation to *GAPDH* housekeeping gene.

**F.** WG32 cells were transfected as in **E** and 48 hours post-transfection cells were treated with 1 μM SB939 or DMSO for 24 hours. Samples were then subjected to western blotting with the antibodies indicated. A representative experiment from three independent repeats is shown. A bottom panel shows quantitation of H3.3K27M densitometry normalised to β-actin.

**Supplementary Tables**

**Supplementary Table 1. Cell lines**

| Cell line | Diagnosis | Brain location | H3 status | Reference or source |
|---|---|---|---|---|
| WG27 | DMG | Pons | H3.1K27M *(HIST1H3B* | Derived from biopsy taken at the Children's Memorial Health Institute (CMHI) in Warsaw |
| WG30 | DMG | Thalamus | H3.3K27M *(H3F3A)* | Derived from biopsy taken at the CMHI |
| WG32 | DMG | Pons | H3.3K27M *(H3F3A)* | Derived from biopsy taken at the CMHI |
| SF8628 | DIPG | Pons | H3.3K27M *(H3F3A)* | Mueller et al., Neuro Oncol 2014 |
| SF7761 | DIPG | Pons | H3.3K27M *(H3F3A)* | Hashizume et al., 2012, J Neurooncol |
| SU-DIPGXIII parental SU-DIPGXIII-KO$^{H3.3K27M}$ | DIPG | Pons | H3.3K27M *(H3F3A)* | Krug et al., Cancer Cell, 2019 |
| BT245 | GBM | Thalamus | H3.3K27M *(H3F3A)* | Krug et al., Cancer Cell, 2019 |
| HSJ019 | GBM | Thalamus | H3.3K27M *(H3F3A)* | Harutyunyan et al., Cell Reports 2020 |
| 7316-1769 | HGG | Thalamus | H3.3K27M *(H3F3A)* | Children's Brain Tumor Tissue Consortium (CBTTC), The Children's Hospital of Philadelphia (CHOP) |
| 7316-1763 | HGG | Thalamus | H3.3K27M *(H3F3A)* | CBTTC, CHOP |
| 7316-195 | HGG | Cerebellum | H3.3K27M *(H3F3A)* | CBTTC, CHOP |
| 7316-913 | HGG | Temporal Lobe | H3 wild-type | CBTTC, CHOP |
| 7316-1746 | HGG | Cerebellum | H3 wild-type | CBTTC, CHOP |

| | | | | |
|---|---|---|---|---|
| 7316-85 | HGG | Temporal Lobe | H3 wild-type | CBTTC, CHOP |

**Supplementary Table 2. siRNA sequences**

| siRNA name | siRNA sequence or purchase information (sense 5’-3’) |
|---|---|
| NTC-control | CUCUCGCUUGGGCGAGAGUAAGtt |
| H2AFZ-si1 | CCGUAUUCAUCGACACCUAtt |
| H2AFZ-si2 | GACUUAAAGGUAAAGCGUAtt |
| H2AFV-si1 | GCAGUAUCUCGCUCACAGAtt |
| H2AFV-si2 | CAGGUAAUGCUUCUAAGGAUCUCAA |
| HDAC1-si | ON-TARGETplus Human HDAC1 (3065) siRNA, L-003493-00-0020 (Dharmacon) |
| HDAC2-si | ON-TARGETplus Human HDAC2(3066)siRNA, L-003495-02-0020 (Dharmacon) |

**Supplementary Table 3. RT-PCR primers**

| Gene | Forward primer (5’- 3’) | Reverse primer (5’- 3’) |
|---|---|---|
| *GAPDH* | TCCTGGAACAGCAAAACAAG | CAGCCTCAGGTTGGTTTCAT |
| *H3F3A* | ATTCGCAAACTTCCCTTC | TGCTAGCTGGATGTCTTTT |
| *H3F3B* | CACACCAGCATCATCTTAAC | ACCAACCACACAACACAA |
| *H2AFZ* | TCCAGTGTTGGTGATTCCAG | GCAGAAATTTGGTTGGTTGG |
| *H2AFV* | TCCCTCACATCCACAAATCTC | AGTACAATGACGGGGAGGAA |

*Supplementary Figure 1*

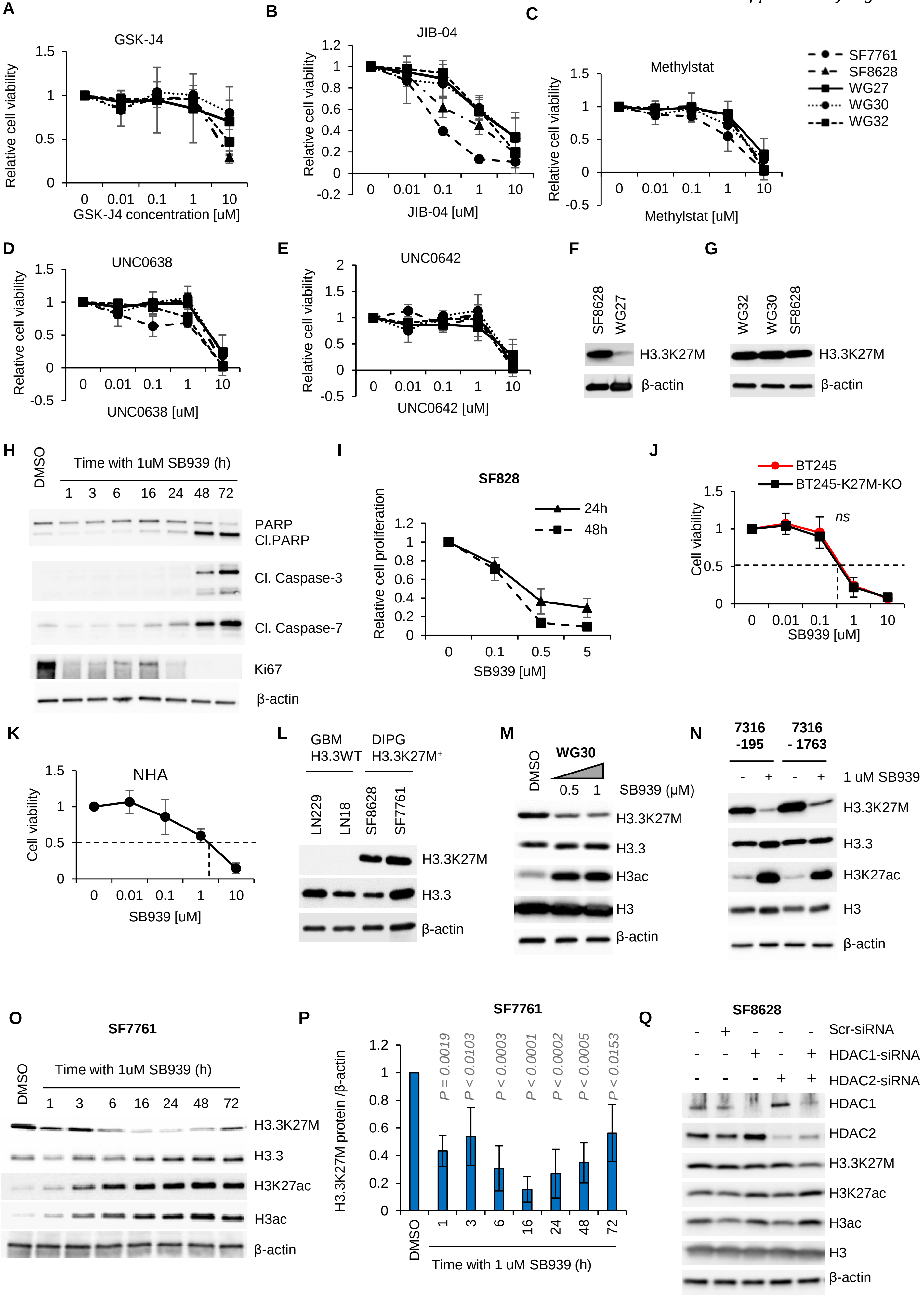

*Supplementary Figure 2*

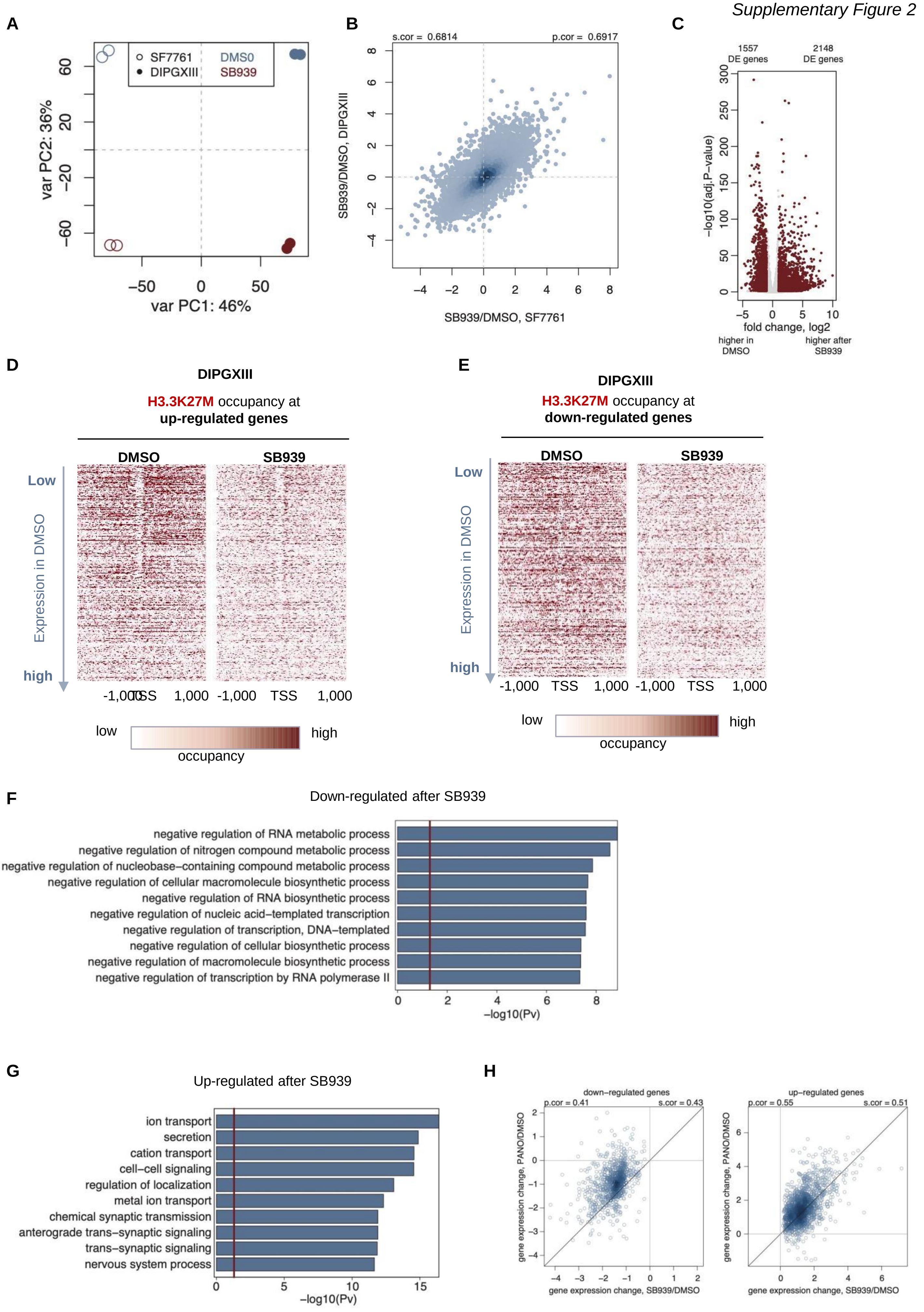

**A**

Down-regulated geness with H3.3K27M loss after SB939

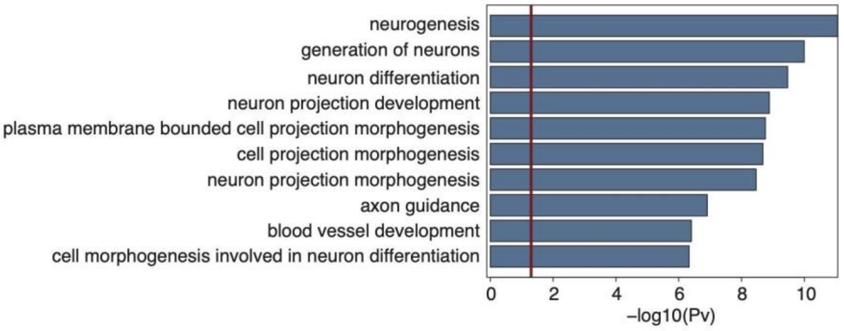


**B**

DIPGXIII

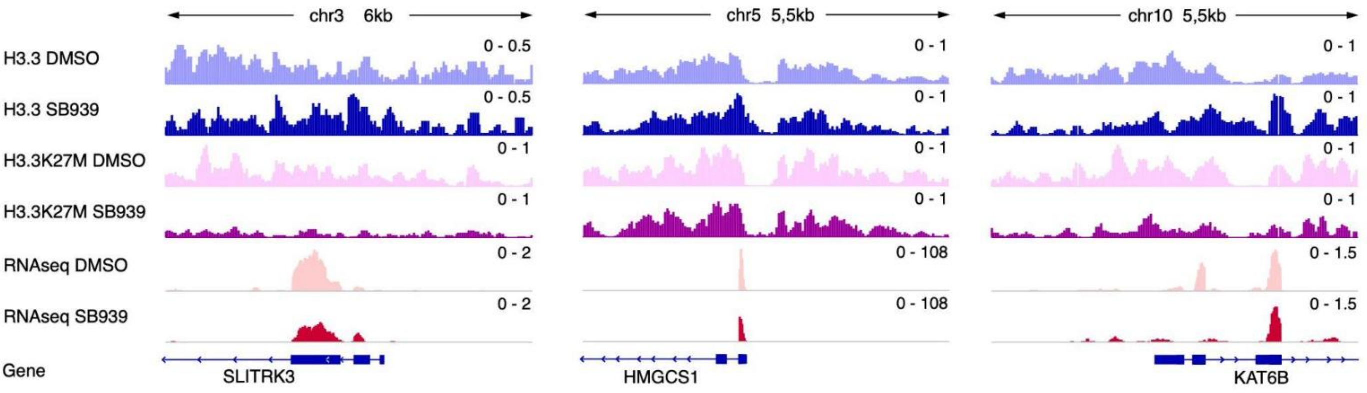


**C**

Expression of all 143 H3K27M-dependent genes from Bender *et al.*. after SB939 treatment

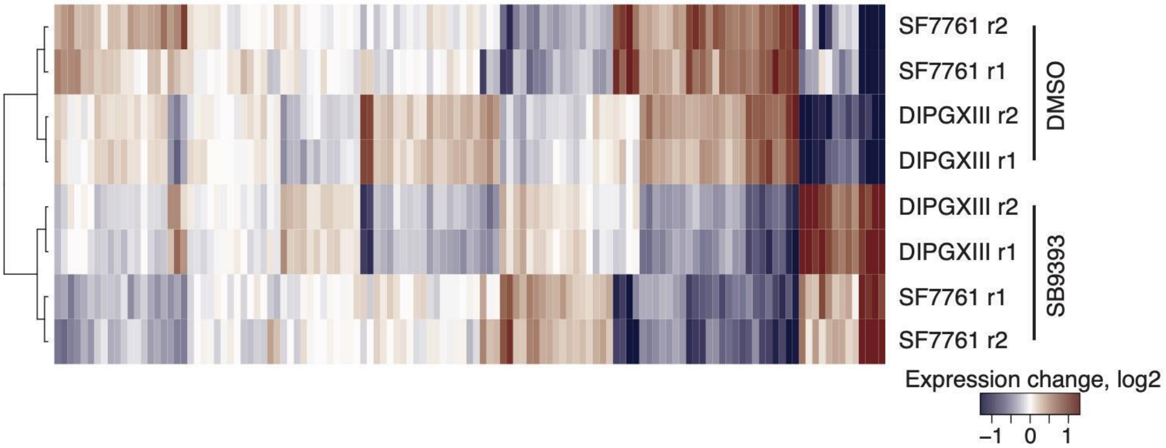

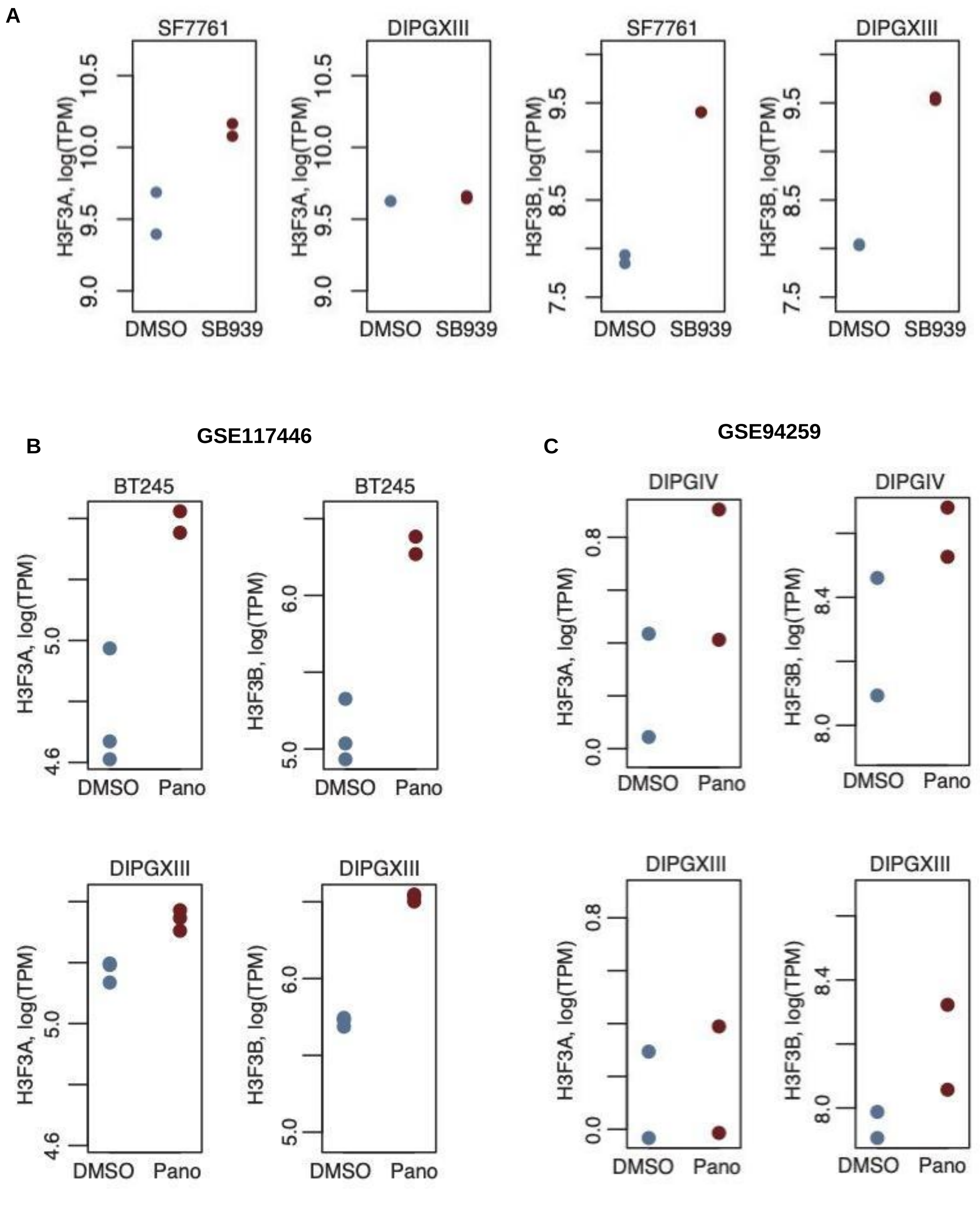
A
SF7761
DIPGXIII
SF7761
DIPGXIII
H3F3A, log(TPM)
H3F3B, log(TPM)
9.0
9.5
10.0
10.5
7.5
8.5
9.5
DMSO
SB939
B
GSE117446
BT245
DIPGXIII
H3F3A, log(TPM)
H3F3B, log(TPM)
4.6
5.0
6.0
DMSO
Pano
C
GSE94259
DIPGIV
DIPGXIII
H3F3A, log(TPM)
H3F3B, log(TPM)
0.0
0.8
8.0
8.4
DMSO
Pano

Supplementary Figure 5

A

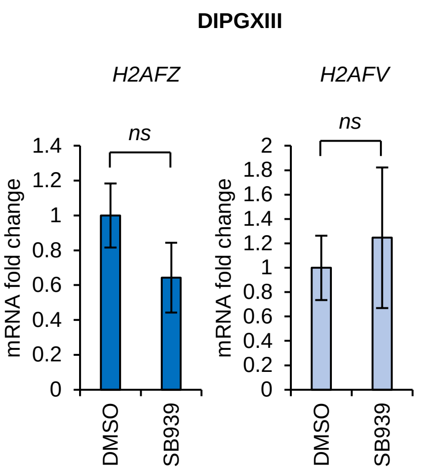


B

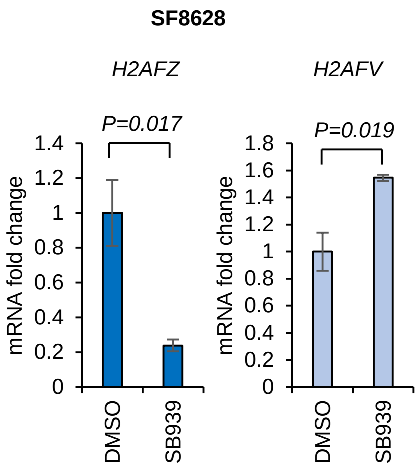


C

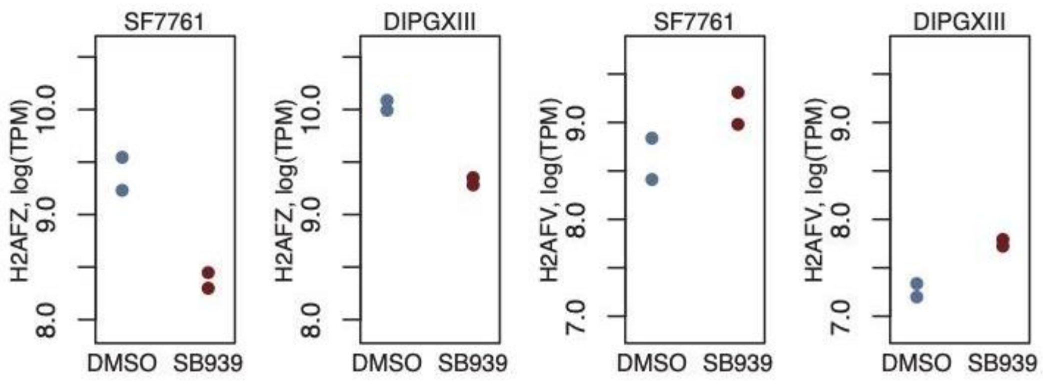


D

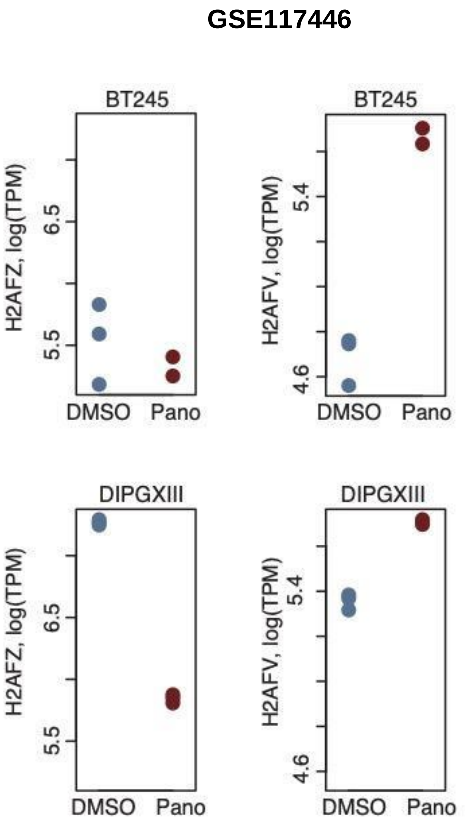


E

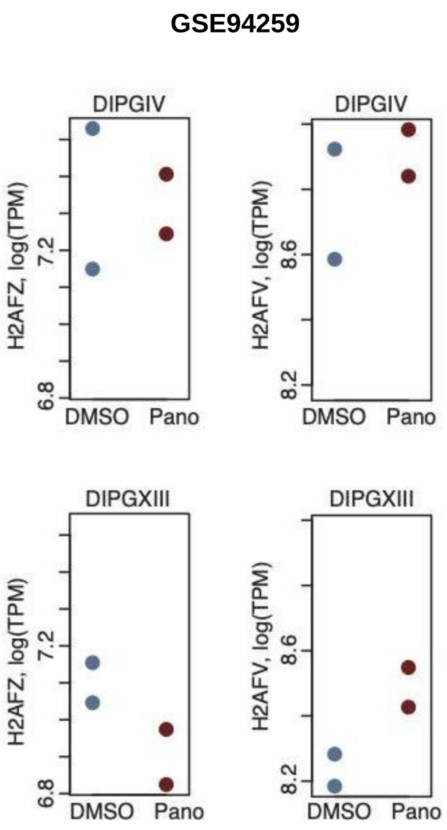

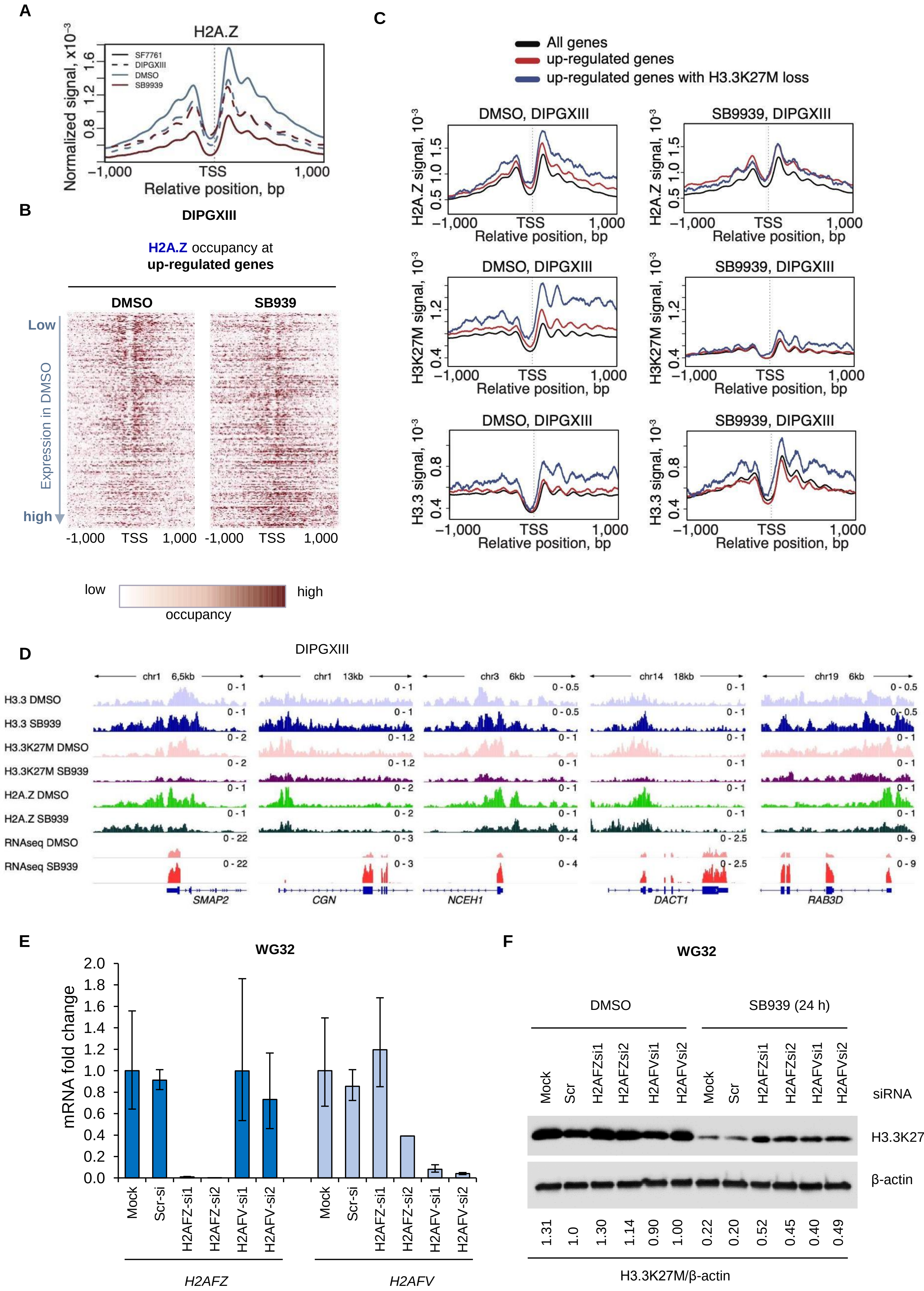

A
H2A.Z
SF7761
DIPGXIII
DMSO
SB9939
Normalized signal, x10⁻³
Relative position, bp
B
DIPGXIII
H2A.Z occupancy at
up-regulated genes
DMSO
SB939
Low
Expression in DMSO
high
low
high
occupancy
C
All genes
up-regulated genes
up-regulated genes with H3.3K27M loss
DMSO, DIPGXIII
SB9939, DIPGXIII
H2A.Z signal, 10⁻³
H3K27M signal, 10⁻³
H3.3 signal, 10⁻³
Relative position, bp
D
DIPGXIII
H3.3 DMSO
H3.3 SB939
H3.3K27M DMSO
H3.3K27M SB939
H2A.Z DMSO
H2A.Z SB939
RNAseq DMSO
RNAseq SB939
SMAP2
CGN
NCEH1
DACT1
RAB3D
E
WG32
mRNA fold change
Mock
Scr-si
H2AFZ-si1
H2AFZ-si2
H2AFV-si1
H2AFV-si2
H2AFZ
H2AFV
F
WG32
DMSO
SB939 (24 h)
siRNA
H3.3K27
β-actin
H3.3K27M/β-actin
